# Ribosome Quality Control Mechanism Mitigates the Cytotoxic Impacts of Ribosome Collisions Induced by 5-Fluorouracil

**DOI:** 10.1101/2023.12.26.573247

**Authors:** Susanta Chatterjee, Parisa Naeli, Nicole Simms, Aitor Garzia, Angela Hackett, Kelsey Coyle, Patric Harris Snell, Tom McGirr, Tanvi Nitin Sawant, Kexin Dang, Zornitsa V. Stoichkova, Tommy Alain, Thomas Tuschl, Simon S. McDade, Daniel B. Longley, Christos G. Gkogkas, Colin Adrain, John R.P. Knight, Seyed Mehdi Jafarnejad

## Abstract

Translation of aberrant or damaged mRNAs results in ribosome stalling and collisions. The Ribosome Quality Control (RQC) mechanism detects collided ribosomes and removes aberrant mRNAs and nascent peptides, thus preventing their cytotoxic effects. Conversely, excessive or unresolved ribosome collisions can induce apoptosis. 5-Fluorouracil (5FU) forms the backbone of standard-of-care chemotherapeutic regimens for several types of cancer. Although best known for its incorporation into DNA and inhibition of thymidylate synthase, a major determinant of 5FU’s anticancer activity is its incorporation into RNAs. Nevertheless, the mechanism(s) underlying RNA-dependent 5FU cytotoxicity and the cellular response to its impact on RNA metabolism remain unclear. Here, we report a key role for RQC in mitigating the cytotoxic effects of 5FU-induced dysregulation of mRNA translation. We show that acute 5FU treatment results in the rapid induction of the mTOR signalling pathway, an enhanced rate of mRNA translation initiation, and increased ribosome collisions that trigger RQC. We also found that RQC deficiency, caused by the depletion of ZNF598, results in increased 5FU-induced cell death, a phenotype that is reversed by inhibition of mTOR or repression of mRNA translation initiation. Importantly, 5FU treatment enhances the expression of key RQC factors, including ZNF598 and GIGYF2, via an mTOR-dependent post-translational regulation mechanism. This acute adaptation likely mitigates the cytotoxic consequences of increased ribosome collisions upon 5FU treatment. Overall, our data indicate a heretofore unknown mTOR-dependent mechanism that augments the RQC process, mitigating the cytotoxicity of 5FU and undermining its anticancer efficacy.

## Introduction

The optimal use of many anti-cancer treatments is impeded by an insufficient understanding of their mechanisms of action. 5-fluorouracil (5FU) is the most commonly used chemotherapeutic agent and backbone of standard-of-care chemotherapy regimens for several types of solid tumours, including colorectal, pancreatic, breast, and head and neck cancers. 5FU is a pyrimidine analogue that, once in the cell, is converted to 3 active metabolites: i) FdUMP, which inhibits the nucleotide synthetic enzyme thymidylate synthase (TS); ii) FdUTP, which mis-incorporates into DNA; and iii) FUTP that mis-incorporates into RNAs^1^. Misincorporation of FUTP into RNAs is a major component of the anticancer activity of 5FU, as RNAs accumulate ∼3000-15000x more 5FU metabolites than DNA^1–7^. This results in the production of abnormal RNAs (e.g., through altered splicing)^2^, aberrant post-transcriptional modifications (e.g. pseudo-uridylation)^7^, alteration of ribosome biogenesis or functions^8^, and aberrant translation-dependent events (e.g., stop-codon readthrough)^4,8^. However, the underlying mechanistic details of RNA-dependent 5FU cytotoxicity remains ill-defined^9^.

The eukaryotic mRNA translation process consists of three main stages: initiation, elongation, and “termination and recycling of the ribosomes”. Initiation is facilitated by a 5’ m7GpppN cap structure and the aid of the eukaryotic Initiation Factor 4F (eIF4F) complex, which consists of the cap-binding subunit eIF4E, RNA helicase eIF4A, and the large subunit eIF4G. eIF4F recruits the 43S pre-initiation complex (PIC) via eIF3. This results in the formation of the 48S PIC that scans the 5’ UTR until a start codon is recognised^10^. Upon detection of the start codon by the PIC, the 60S ribosomal subunit joins to form the 80S ribosome and initiates the “elongation phase”, which continues until the ribosome arrives at a stop codon, whereupon translation termination can occur^11^. Initiation is the main stage of regulation of translation, which is achieved via several mechanisms [reviewed in^12^], including abrogation of the interaction between eIF4E and eIF4G by the family of small eIF4E-binding proteins (4E-BPs). Affinity of 4E-BPs for eIF4E is regulated by phosphorylation of 4E-BPs on several residues by the mechanistic Target of Rapamycin Complex 1 (mTORC1) that controls the rate of translation initiation and other key metabolic processes in response to external and internal stimuli, such as nutrient availability^13^.

Beyond the imperative to regulate translation rate during initiation^10^, the synthesis of full-length, correctly folded proteins is vital for maintaining homeostasis and preventing diseases. Translating ribosomes can be stalled by obstacles such as the presence of damaged (e.g., oxidized) or modified nucleotides and premature poly(A) sequence within the coding region^14–19^. Notably, efficiently translated mRNAs are more prone to ribosome collisions^20–22^, due to the higher chance of presence of trailing ribosomes that collide into a stalled ribosome. Unresolved stalled ribosomes could have deleterious consequences due to protein aggregation or release of truncated proteins. The Ribosome Quality Control (RQC) mechanism detects collided ribosomes and removes the aberrant mRNAs and nascent peptides, thus preventing their cytotoxic consequences^23,24^. Conversely, an overwhelmed (e.g. due to excessive mRNA damage) or dysfunctional RQC results in cell death^25^.

Significant progress has been made in understanding the molecular mechanisms of RQC^26^, including identification of crucial RQC factors, such as the RNA-binding and E3 ubiquitin ligase protein ZNF598, which triggers RQC by promoting mono-ubiquitination of the small ribosomal subunits^19,27,28^. ZNF598, also recruits the cap-binding protein and translational repressor 4EHP via GIGYF2 and thereby prevents further translation initiation on the aberrant mRNA^29^. Alternatively, EDF1 can act as a sensor of ribosome collisions and recruit GIGYF2-4EHP complex at collided ribosomes, independent of ZNF598^30^. The ASCC complex containing ASCC3, ASCC2, and TRIP4, participates in the disassembly of colliding ribosomes. Subsequently, PELO and ABCE1 split the ribosomal subunits, and ANKZF1/VMS1, in cooperation with p97 and Arb1, release the nascent peptide chain from 60S complexes. In parallel, the Ltn1/VCP/NEMF complex promotes the proteasomal degradation of the nascent peptide^26^. Through this process, RQC protects the integrity of mRNA translation and contributes to the maintenance of cellular homeostasis^31,32^. Conversely, defects in RQC, for instance due to depletion of Ltn1 or NEMF, lead to major pathological consequences such as neurological disorders resulting from accumulation of toxic protein aggregates^15,24,33,34^. Nevertheless, while the causal implication of the general mRNA translation machinery in tumorigenesis and therapy resistance is well-documented^12,35^, little is known about the role of RQC in cellular responses to anti-cancer treatments that impact RNA metabolism.

Here, we demonstrate that, contrary to previous assumptions, acute 5FU treatment induces a major reprogramming of the cellular mRNA translation machinery that includes activation of the mTORC1 signalling pathway, an elevated rate of translation initiation, and increased ribosome collisions. Crucially, we show that RQC mitigates against 5FU-induced toxicity by resolving the collided ribosomes. Conversely, these collided ribosomes accumulate in the RQC-deficient cells, resulting in enhanced 5FU-induced cell death. We also demonstrate that, inhibition of mTOR or repression of cap-dependent mRNA translation reverses the sensitisation to 5FU seen in RQC-deficient cells. Furthermore, our data suggest the presence of a hitherto unknown intrinsic cellular mechanism of post-translational upregulation of expression of the key RQC factors ZNF598 and GIGYF2 upon mTOR activation. Thus, 5FU treatment leads to an mTOR-dependent upregulation of these RQC factors, which further bolsters the cellular response to the 5FU-induced ribosome collisions and mitigates against 5FU-induced cytotoxicity.

## Results

### Acute 5FU treatment enhances global mRNA translation

Recent studies suggest that 5FU treatment results in downregulation of mRNA translation^4,8,36^. However, these studies predominantly focussed on the impact on mRNA translation following 5FU exposures of longer than 24 hours. We postulated that such analyses might be susceptible to biases due to the cytotoxic effects of 5FU, including stress induced by DNA damage upon prolonged 5FU treatment. Consequently, these analyses may not accurately reflect the effect of 5FU on mRNA translation.

Using an antibody able to detect 5-fluorouridine incorporated into nucleic acids^37,38^, we assessed the covalent immobilisation of 5FU metabolites within single cells. We observed a rapid cytoplasmic 5FU localisation within 30 min of treatment, which remained at similar levels up to 24 h (**Fig. 1A**) and was very sensitive to RNase digestion (**Supp. Fig. 1A**). These data emphasize that 5FU has a substantial and early impact on RNA metabolism and underscore the importance of examining the impact of 5FU on mRNA translation during these early timepoints. Therefore, we sought to investigate the changes in the mRNA translation profile upon acute treatment of HCT116 (colorectal cancer) and SUIT-2 (pancreatic adenocarcinoma) cells with 5FU. Surface Sensing of Translation (SUnSET) assay, a nonradioactive puromycin/antibody-based tool for quantification of protein synthesis^39^, revealed that while prolonged (48 h) 5FU treatment slightly reduced the rate of new protein synthesis in HCT116 and SUIT-2 cells, shorter 5FU treatment (<12 h) surprisingly resulted in increased rate of protein synthesis (**Fig. 1B** **& Supp. Fig. 1B**). The enhanced rate of protein synthesis upon acute 5FU treatment was corroborated by a pulse-labelling assay with the methionine analogue L-homopropargylglycine^40^ (**Fig. 1C** **& Supp. Fig. 1C**). Furthermore, polysome profiling assay to analyse the association (loading) of mRNAs with ribosomes revealed a shift from the untranslated or weakly translated sub-polysome region to polysomes upon acute 5FU treatment (**Fig. 1D**), which was reversed at 48 h post-treatment (**Supp. Fig. 1D**). This indicates that mRNA-ribosomes association is enhanced upon acute 5FU treatment.

**Figure 1:**
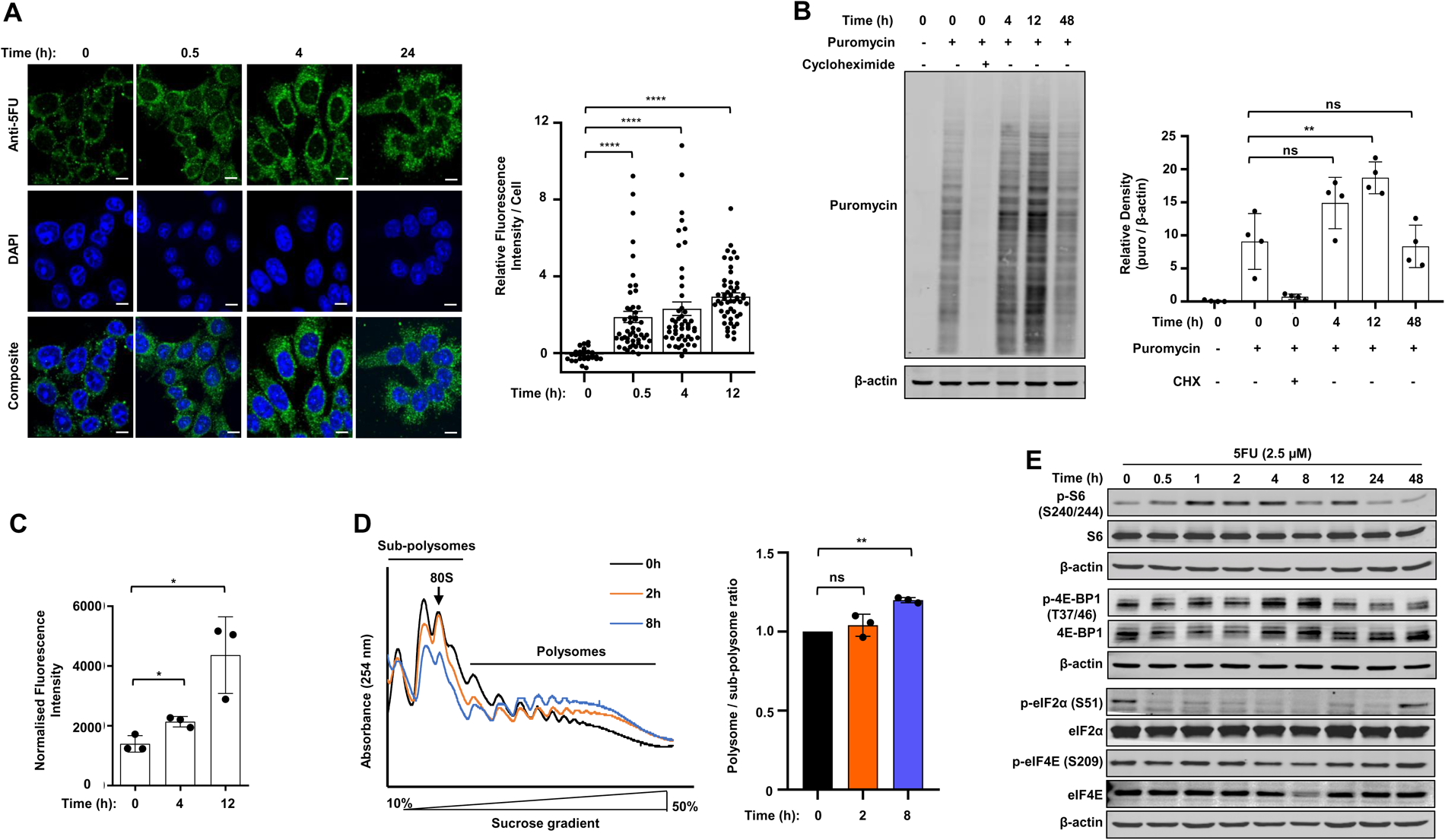
Increased mRNA translation upon acute 5FU treatment. (**A**) *Left*: HCT116 Cells were treated with 5FU (2.5 µM) for 0.5, 4, and 24 h for immunofluorescence analysis of 5FU incorporation. Cells were stained for 5-FU (green) using anti-BrdU antibody and counterstained with the nucleic acid dye DAPI (blue). An untreated control (0 h) was included, which displays low level of background. *Right:* Graph showing 5FU fluorescence per cell (29 cells for the 0 h and 50 cells quantified in the 0.5 h, 4 h, and 24 h groups) relative to 0 h control. **** p<0.0001, one way ANOVA with Dunnett’s multiple comparisons test. Scale bar = 45 µm. (**B**) Quantification of new protein synthesis by Surface Sensing of Translation (SUnSET) assay in HCT116 cells treated with 5FU (2.5 µM) for the indicated times. *Left*: Representatives immunoblot analysis of lysates probed with the indicated antibodies. *Right*: The bar graph represents the relative change in Puromycin / β-actin signal density in each group, measured by ImageJ. Data are presented as mean ± SD; n=4 independent replicates; **p<0.01, two-tailed student’s t-test. (**C**) Quantification of new protein synthesis by pulse labelling with the methionine analogue L-homopropargylglycine (HPG). HCT116 cells were treated with 5FU (2.5 µM) for the indicated times. The signal was normalized against the unstained samples for each group. Data are presented as mean ± SD; n=3 independent replicates; *p<0.05, two-tailed student’s t-test. (**D**) Analysis of general mRNA-ribosome association in HCT116 cells. *Left.* Cells were treated with 5FU (2.5 µM) for the indicated times before harvesting and analysis using polysome profiling assay. A shift from the sub-polysome to polysomes fractions indicates enhanced mRNAs-ribosomes association. *Right.* Quantification of the polysome/sub-polysome ratio in control and 5FU treated cells. Data are presented as mean ± SD; n=3 independent replicates; **p< 0.01, two-tailed student’s t-test. (**E**) Western blot analysis of effects of 5FU treatment (2.5 µM) for the indicated times on the signalling pathways regulating mRNA translation machinery in HCT116 cells. To avoid excessive non-specific signals due to repeated re-probing, identical samples were run on separate gels. β-actin was used as loading control for each membrane. Quantification of the pS6 and p4E-BP1 expression is presented in Supp. Fig. 1E & F. *Also see the related* Supp. Figure 1.

We next investigated the impacts of acute and prolonged 5FU treatment on core mechanisms of regulation of mRNA translation initiation. Acute 5FU treatment markedly and rapidly (<30 min) induced phosphorylation of RPS6 (S240/244) and 4E-BP1 (T37/46), key markers of mTORC1 activity, which decreased with longer (>12 h) treatment (**Fig. 1E** **& Supp. Fig. 1E-G**). Notably, acute 5FU treatment did not activate the Integrated Stress Response (ISR) pathway (indicated by eIF2α phosphorylation on S51 residue) or the MNK-regulated eIF4E phosphorylation on S209 residue (**Fig. 1E** **& Supp. Fig. 1G**). Altogether, these data suggest that acute 5FU-treatment enhances global mRNA translation via activation of the mTORC1 pathway.

### RQC resolves the 5FU-induced ribosome collisions

We hypothesised that the enhanced rate of mRNA translation (**Fig. 1**), combined with the known impacts of 5FU on production of abnormal RNAs^2^, post-transcriptional modifications^7^, alteration of ribosome functions^8^, and aberrant translation events (e.g. stop-codon readthrough^4,8^) could lead to ribosome collisions and triggering of the RQC mechanism. We observed an increased mono-ubiquitination of ribosomal small subunit protein eS10, a marker of ribosome collision and activation of RQC^19^, upon 5FU treatment (**Fig. 2A**). The RNA-binding protein and E3 ubiquitin ligase ZNF598 plays a key role in detection of ribosome collisions and triggering RQC, and its deletion abrogates RQC^19,27,28^. To investigate the role of RQC in the cellular response to 5FU, we generated ZNF598-knockout (KO) HCT116 and SUIT-2 cells with CRISPR-Cas9 (**Supp. Fig. 2A**). We observed that unlike in the parental cells, where 5-FU treatment triggered eS10 mono-ubiquitination, this effect was absent in ZNF598-KO cells (**Fig. 2B** **& Supp. Fig. 2B**). Considering the observed increased general mRNA translation and protein synthesis upon acute 5FU treatment (**Fig. 1B-D**), we next assessed the impact of ZNF598-KO on 5FU-induced mRNA translation. Polysome profiling revealed that similar to the parental cells (**Fig. 1D**) acute 5FU treatment increased mRNA-ribosome association in ZNF598-KO HCT116 cells (**Fig. 2C**). However, pulse-labelling with the methionine analogue L-homopropargylglycine for analysis of nascent protein synthesis showed that compared with the parental cells, ZNF598-KO diminished the acute 5FU-induced protein synthesis (**Fig. 2D**). We hypothesized that this discrepancy in 5FU-induced mRNA-ribosome association (detected via polysome profiling) versus protein synthesis (observed in pulse labelling) in ZNF598-KO cells is due to the inability of the RQC-deficient cells to resolve 5FU-induced ribosome collisions. This deficiency results in accumulation of non-productive stalled ribosomes in polysomes and reduction of protein synthesis in ZNF598-KO cells. To test this, we assessed the occurrence of ribosome collisions by Micrococcal Nuclease (MNase) digestion assay in 5FU-treated parental and ZNF598-KO cells. MNase degrades the mRNA between ribosomes, thereby releasing 80S ribosomes. However, when ribosome collisions occur, the inter-ribosomal mRNA of the collided ribosomes (disome and trisomes) is protected from MNase digestion^41^. We observed that while the untreated ZNF598-KO cells accumulate only a small amount of disomes compared with their parental counterparts (**Fig. 2E**), 5FU treatment results in a clear accumulation of disome and trisome peaks in ZNF598-KO but not in parental cells (**Fig. 2F**). These data further indicate an enhanced rate of ribosome collisions upon 5FU treatment that are readily resolved by RQC mechanism in parental cells, whereas RQC-deficient ZNF598-KO cells are impaired in their ability to resolve the accumulated collided ribosomes.

**Figure 2:**
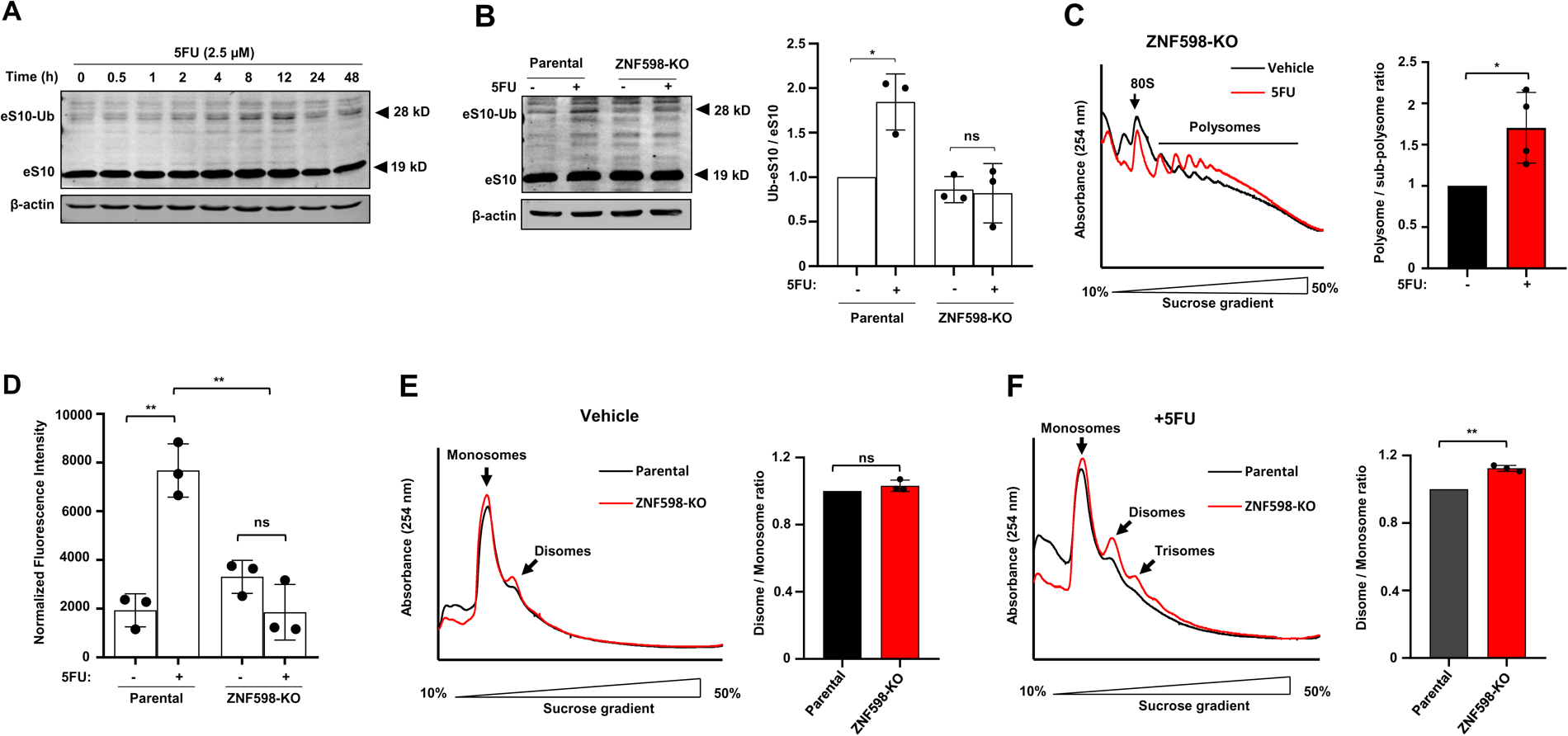
ZNF598 resolves 5FU induced collided ribosomes. (**A**) Western blot analysis of mono-ubiquitination of eS10 upon 5FU treatment (2.5 µM) for the indicated time in HCT116 cells. β-actin was used as a loading control. (**B**) *Left.* Western blot analysis of mono-ubiquitination of eS10 in parental and ZNF598-KO HCT116 cells upon 5FU treatment (2.5 µM) for 1 h. β-actin was used as loading control. *Right:* Densitometric quantitation of the mono-ubiquitinated eS10 / total eS10. Data are normalised against the respective control for each replicate and presented as mean ± SD; n=3 independent replicates; *p< 0.05, two-tailed student’s t-test. (**C**) *Left*: Polysome profiling analysis of general mRNA-ribosome association in in ZNF598-KO HCT116 cells treated with 5FU (2.5 µM) for 6 h or vehicle using polysome profiling assay. *Right*: Quantification of the polysome/sub-polysome ratio in vehicle and 5FU treated cells. Data are normalised against the respective control for each replicate and presented as mean ± SD; n=4 independent replicates; *p<0.05, two-tailed student’s t-test. (**D**) Quantification of new protein synthesis by pulse labelling with the methionine analogue L-homopropargylglycine in parental and ZNF598-KO HCT116 cells treated with 5FU (2.5 µM) for 4 h. Data are normalised against the respective control for each replicate and presented as mean ± SD; n=3 independent replicates; **p<0.01, two-tailed student’s t-test. (**E**) *Left:* Assessment of ribosome collisions in parental and ZNF598-KO HCT116 cells by polysome profiling following micrococcal nuclease (MNase) digestion of cell lysates. *Right:* Quantification of the disome/monosome ratio in each condition. Data are normalised against the respective control for each replicate and presented as mean ± SD; n=3 independent replicates; ns=non-significant; two-tailed student’s t-test. (**F**) *Left:* Micrococcal nuclease (MNase) digestion in parental and ZNF598-KO HCT116 cells treated with 5FU (2.5 µM) for 12 h. *Right:* Quantification of the disome/monosome ratio in each condition. Data are normalised against the respective control for each replicate and presented as mean ± SD; n=3 independent replicates; **p<0.01, two-tailed student’s t-test. *Also see the related* Supp. Figure 2.

### RQC deficiency enhances the sensitivity of cancer cells to 5FU

Based on our findings, we hypothesised that the ability of the cells to resolve 5FU-induced collided ribosomes by RQC may mitigate against 5FU-induced lethality in cancer cells. Our initial characterisation of the ZNF598-KO HCT116 and SUIT-2 cells revealed that ZNF598-KO resulted in a slightly reduced growth rate (**Fig. 3A** **& Supp. Fig. 3A**). Importantly, ZNF598-KO markedly increased the 5FU sensitivity of HCT116 (IC_50_=0.73 µM and 0.19 µM in parental and ZNF598-KO, respectively; **Fig. 3B**) and SUIT-2 cells (IC_50_=1.36 µM and 0.23 µM in parental and ZNF598-KO respectively; **Supp. Fig. 3B**). Similar results were observed upon depletion of ZNF598 expression by three different shRNAs in HCT116 cells (**Supp. Fig. 3C**). Enhanced 5FU-induced toxicity upon ZNF598-KO was also corroborated by impaired clonogenic survival (**Fig. 3C** **& Supp. Fig. 3D**). Moreover, these enhanced growth inhibitory effects correlated with elevated 5FU-induced apoptosis, assessed by PARP cleavage in ZNF598-KO cells (**Fig. 3D** **& Supp. Fig. 3E**). Previous reports suggested that increased levels of unresolved collided ribosomes in ZNF598-KO cells RQC leads to activation of the Ribotoxic Stress Response (RSR) and apoptosis via ZAKa/p38 pathway^25^. We noted that while acute 5FU treatment did not induce p38 phosphorylation (a marker of RSR activation) in parental cells, RQC-deficient ZNF598-KO cells exhibited distinctively increased p38 phosphorylation upon 5FU treatment (**Fig. 3E**). These data strongly suggest a cytoprotective role for ZNF598/RQC in cellular response to 5FU, creating a dependency on RQC to mitigate the cytotoxic impacts of 5FU. To assess the importance of misincorporation of FdUTP into DNA and FdUMP-induced thymidylate synthase (TS) inhibition for increased sensitivity of ZNF598-KO cells to 5FU, we compared the effects of ZNF598-KO on the cell viability in response to treatment with FUDR, the direct precursor for FdUMP and FdUTP, and FUTP (which mis-incorporates into RNAs). ZNF598-KO increased sensitivity to FUDR (**Fig. 3F** **& Supp. Fig. 3F**) and to an even greater extent FUTP (**Fig. 3G** **& Supp. Fig. 3G**). In contrast, ZNF598-depletion did not have an impact on cancer cells’ sensitivity to Gemcitabine (**Fig. 3H** **& Supp. Fig. 3H**), another nucleic-acid-incorporating chemotherapeutic reagent^41^. These data indicate that the RQC machinery has a selective role in mitigating the cellular response to 5FU-induced toxicity, which can be uncoupled from its incorporation into DNA or thymidylate synthase inhibition and the mechanisms of action of other RNA-incorporating antimetabolites.

**Figure 3:**
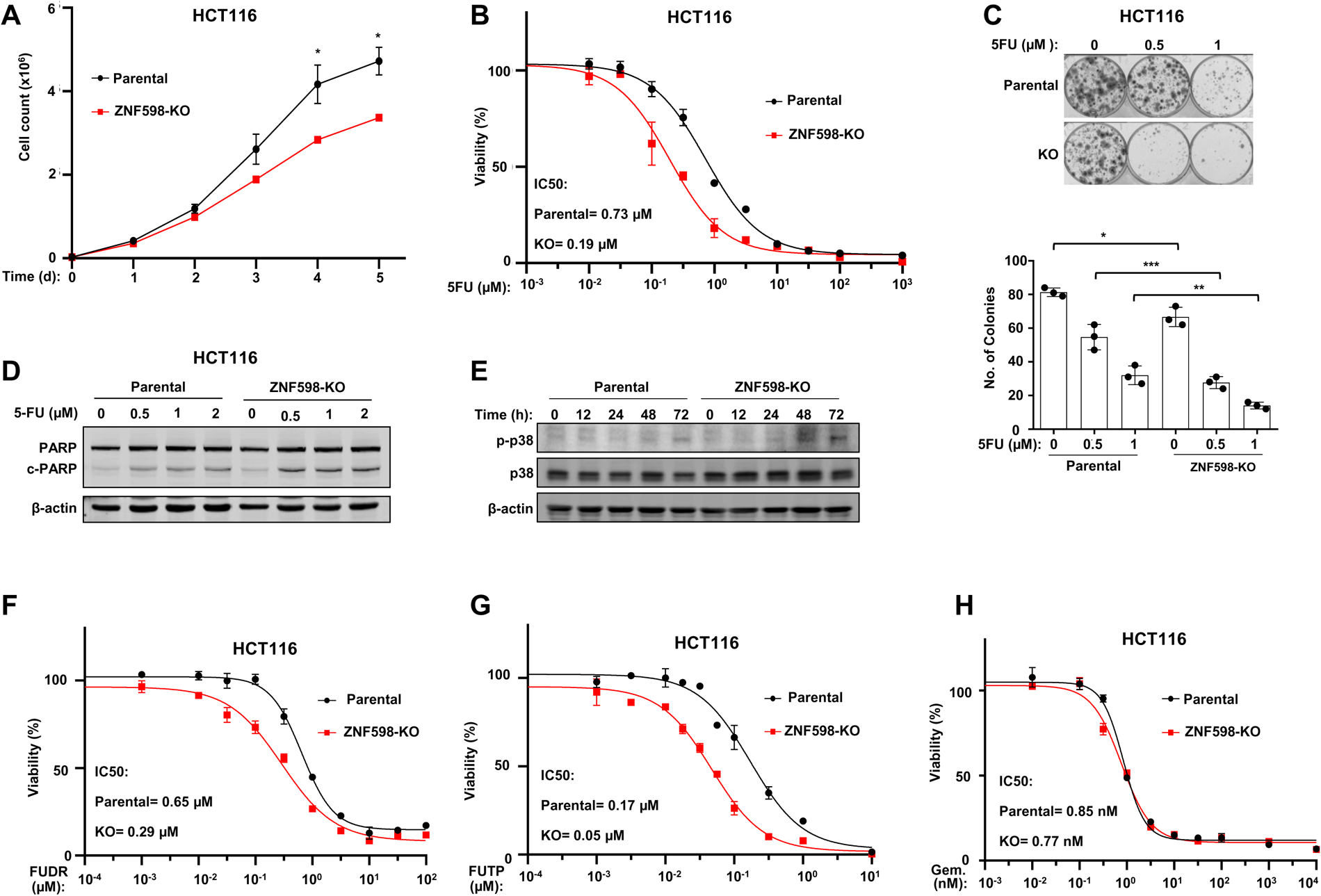
Defective RQC results in increased sensitivity of cancer cells to 5FU. (**A**) Cell growth assay with parental and ZNF598-KO HCT116 cells after the indicated time points. Data are shown as mean ± SD; n=3 independent replicates; *p<0.05, two-tailed student’s t-test. (**B**) Dose-response assay for measurement of sensitivity of parental and ZNF598-KO HCT116 cells to 5FU. Cell viability was measured 72 h post-treatment using CellTiter-Glo® Luminescent Cell Viability Assay. (**C**) *Top*: Colony formation assay with parental and ZNF598-KO HCT116 cells 12 days post-seeding. Cells were treated with the indicated concentration of 5FU. *Bottom*: Quantification of number of colonies with a diameter >200 µm in each well. Data are presented as mean ± SD; n=3 independent replicates; *p<0.05; **p<0.01; ***p<0.001, two-tailed student’s t-test. (**D**) Western blot analysis of expression of cleaved PARP (c-PARP) using lysates derived from parental and ZNF598-KO HCT116 cells treated with the indicated doses of 5FU for 72 h. β-actin was used as loading control. (**E**) Western blot analysis of expression of the marker of the Ribotoxic Stress Response (RSR; p-38) in lysates derived from parental and ZNF598-KO HCT116 cells treated with 5FU (2.5 µM) for the indicated times with the indicated antibodies. (**F-H**) Dose-response assay for measurement of sensitivity of parental and ZNF598-KO HCT116 cells to (**F**) FUDR, the precursor of the DNA-incorporating 5FU metabolite, (**G**) RNA-incorporating 5FU metabolite FUTP, and (**H**) Gemcitabine. 72 h post-treatment cell viability was measured using CellTiter-Glo® Luminescent Cell Viability Assay. *Also see the related* Supp. Figure 3.

### Inhibition of translation initiation reduces 5FU-induced ribosome collision and cellular dependency of RQC

We further sought to investigate the underlying mechanism of 5FU-induced ribosomal collision. Our earlier observations (**Fig. 1B-E**) revealed increased mRNA translation initiation and mTORC1 activity upon 5FU treatment. It has been previously reported that increased translation initiation may exacerbate ribosome collisions as it may lead to an increased presence of trailing ribosomes that collide into a stalled ribosome^42–44^. Considering the key role of mTORC1 in regulation of translation initiation^12^, we sought to investigate the impact of blocking mTOR activity on 5FU-induced ribosome collisions. Treatment with the mTOR inhibitor Torin-1, decreased eS10 ubiquitination in the parental HCT116 cells in response to 5FU treatment (**Fig. 4A**) and reduced 5FU-induced accumulation of disomes and trisomes in ZNF598-KO cells (**Fig. 4B**). We reasoned that if 5FU-induced ribosome stalling contributes to cytotoxicity, the mTOR-dependent reduction in ribosome stalling would alter cell fate. Indeed, we observed that mTOR inhibition reverses the sensitisation to 5FU seen in ZNF598-KO cells (**Fig. 4C**). Importantly, mTOR has multiple downstream effectors, which regulate several processes other than mRNA translation^13^. Thus, we sought to use orthogonal means to verify that our observation of reduced 5FU sensitivity in the RQC-deficient cells upon mTOR inhibition is due to repression of translation initiation. We used the eIF4E/eIF4G interaction inhibitor 1 (4EGI-1), which blocks cap-dependent translation initiation by dissociation of the eIF4F complex^45^. Corroborating our findings with mTOR inhibition, 4EGI-1 treatment also significantly reduced the accumulation of collided ribosomes (disomes and trisomes) in 5FU-treated ZNF598-KO cells (**Fig. 4D**) and alleviated the enhanced 5FU sensitivity of the ZNF598-KO cells compared with the parental cells (**Fig. 4E**).

**Figure 4:**
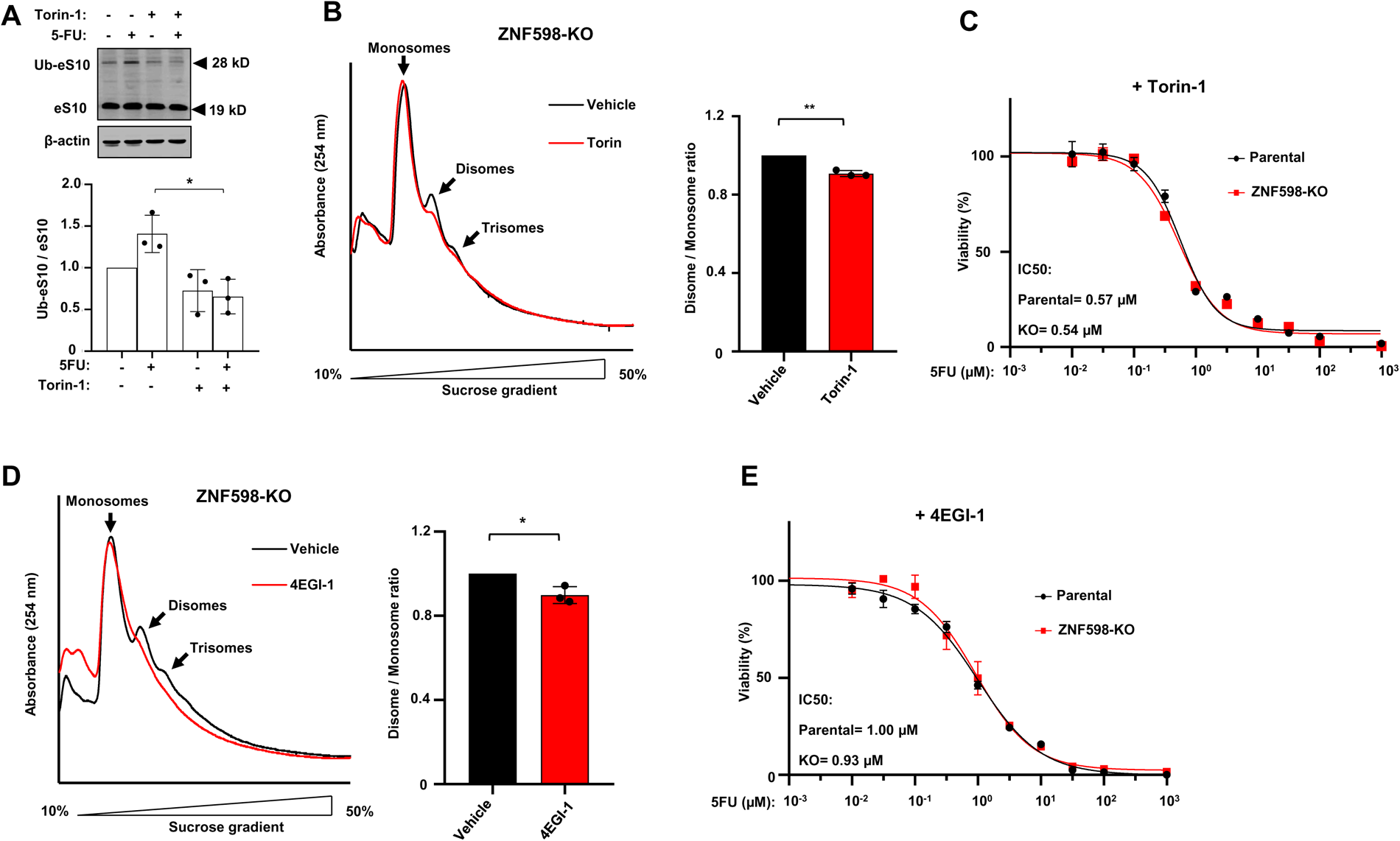
Inhibition of translation initiation reduces the 5FU-induced ribosome collisions and cell death in RQC-deficient cells. (**A**) *Top*: Western blot analysis of mono-ubiquitination of eS10 in lysates derived from HCT116 cells pre-treated with Torin-1 (15 nM) or vehicle for 24 h, followed by treatment with 5FU (2.5 µM) for 1 h. β-actin was used as loading control. *Bottom:* Densitometric quantitation of the mono-ubiquitinated eS10 / total eS10. Data are normalised against the respective control for each replicate and presented as mean ± SD; n=3 independent replicates; *p< 0.05, two-tailed student’s t-test. (**B**) *Left:* Micrococcal nuclease (MNase) digestion in ZNF598-KO HCT116 cells pre-treated with Torin-1 (15 nM) or vehicle for 24 h, followed by treatment with 5FU (2.5 µM) for 12h. *Right:* Quantitation of the disome/monosome ratios. Data are normalised against the respective control for each replicate and presented as mean ± SD; n=3 independent replicates; **p< 0.01, two-tailed student’s t-test. (**C**) Dose-response assay for measurement of 5FU sensitivity of parental and ZNF598-KO HCT116 cells in the presence of Torin-1(15 nM). 72 h post-treatment cell viability was measured using CellTiter-Glo® Luminescent Cell Viability assay. (**D**) *Left.* Micrococcal nuclease (MNase) digestion in ZNF598-KO HCT116 cells pre-treated with 4EGI-1 (25 μM) or vehicle for 24 h, followed by treatment with 5FU (2.5 µM) for 12 h. *Right.* Quantification of the disome/monosome ratio. Data are normalised against the respective control for each replicate and presented as mean ± SD; n=3 independent replicates; *p< 0.05, two-tailed student’s t-test. (**E**) Dose-response assay for measurement of 5FU sensitivity of parental and ZNF598-KO HCT116 cells in the presence of 4EGI-1 (25 μM). 72 h post-treatment cell viability was measured using CellTiter-Glo® Luminescent Cell Viability assay.

Altogether, these data demonstrate that the enhanced rate of mRNA translation initiation upon 5FU treatment contributes to the increased ribosome collisions in 5FU-treated cells and the elevated 5FU-induced cell death in RQC-deficient cells. Thus, blocking mTOR activity or translation initiation reduces the risk of ribosome collisions in 5FU-treated cells and thereby alleviates the requirement for a functional RQC to prevent the cytotoxic consequences of collided ribosomes.

### mTOR-mediated upregulation of key RQC factors by 5FU

Considering the cyto-protective role of ZNF598/RQC against 5FU-treatment, we next sought to investigate the impact of 5FU on expression of ZNF598 and several other factors involved in detection of ribosome collision and triggering RQC, namely EDF1, ASCC3, RACK1, and GIGYF2^26^. Acute 5FU treatment increased the expression of ZNF598 and GIGYF2, a key factor in ZNF598- and EDF1-mediated translational repression of the mRNAs with collided ribosomes^29,30^ and to a lesser extent ASCC3, but not RACK1 or EDF1 (**Fig. 5A** **& Supp. Fig. 4A-C**). Further inspection revealed a remarkably rapid increase in expression of ZNF598 and GIGYF2 within 10 min of 5FU treatment (**Fig. 5B**). RT-qPCR measurement found no significant change in *ZNF598* or *GIGYF2* mRNA levels upon acute 5FU treatment (**Fig. 5C** **& Supp. Fig. 4D**), which suggests involvement of a post-transcriptional mechanism of regulation. To further dissect this mechanism, we assessed the role of translational regulation in increased expression of ZNF598 and GIGYF2. We pre-treated the cells with the translation inhibitor Cycloheximide (CHX), followed by treatment with 5FU. While expression of ZNF598, GIGYF2, and EDF1 was decreased in vehicle-treated cells in the presence of CHX (**Supp. Fig. 5A**), 5FU-treatment increased expression of ZNF598 and GIGYF2, but not EDF1, even in presence of CHX (**Fig. 5D**). These data suggest the presence of a post-translational mechanism of regulation that rapidly increases the expression of the key RQC factors ZNF598 and GIGYF2 in response to 5FU treatment.

**Figure 5:**
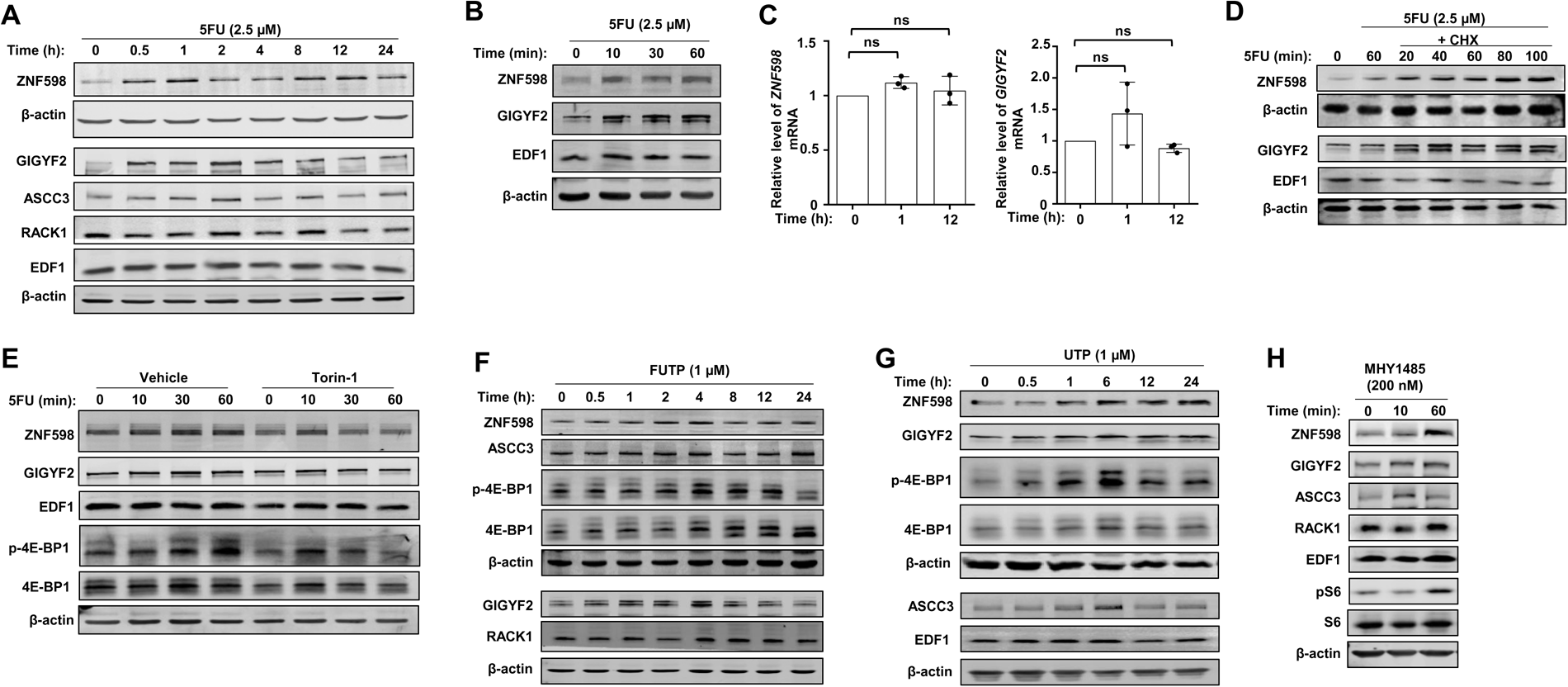
mTOR-dependent upregulation of expression of ZNF598 and GIGYF2 by 5FU-derived metabolites and UTP. (**A**) Western blot analysis of expression of indicated proteins in lysates derived from HCT116 cells treated with 5FU (2.5 µM) for the indicated time points. Quantification of the ZNF598 and GIGYF1 protein expression is presented in Supp. Fig. 4A & B. (**B**) Western blot analysis of expression of the indicated proteins in lysates derived from HCT116 cells treated with 5FU (2.5 µM) for the indicated time points. (**C**) Quantitative RT-PCR analysis of expression of *ZNF598* and *GIGYF2* mRNAs in HCT116 cells upon 5FU treatment (2.5 µM) for the indicated times. Data are presented as mean ± SD; n=3 independent replicates; ns=non-significant; two-tailed student’s t-test. (**D**) Western blot analysis of expression of the indicated proteins in lysates derived from HCT116 cells treated with 5FU (2.5 µM) for the indicated time points in the presence or absence of 100 μg/ml Cycloheximide (CHX). (**E**) Western blot analysis of expression of the indicated proteins in lysates derived from HCT116 cells pre-treated with Torin-1(15 nM) or vehicle for 24 h, followed by treatment with 5FU (2.5 µM) for the indicated time points. Quantification of the ZNF598 and GIGYF1 protein expression is presented in Supp. Fig. 5D & E. (**F & G**) Western blot analysis of lysates derived from HCT116 cells treated with (**F**) FUTP (1 µM) or (**G**) UTP (1 µM) for the indicated times. (**H**) Western blot analysis of lysates derived from HCT116 cells treated with the small molecule mTOR activator MHY1485 (200 nM) for the indicated time points. *Also see the related* Supp. Figure 4 & 5.

Notably, we observed that the rapid upregulation of ZNF598 and GIGYF2 expression upon 5FU treatment coincides with the rapid activation of mTORC1 signalling (**Supp. Fig. 5B**). Therefore, we assessed the possible role of the mTOR pathway in this process. Treatment with mTOR inhibitor Torin-1, resulted in a dose-dependent reduction in ZNF598 and GIGYF2 expression (**Supp. Fig. 5C**). Furthermore, pre-treatment with Torin-1 abrogated the 5FU-induced increase in ZNF598 and GIGYF2 expression (**Fig. 5E, Supp. Fig. 5D & E**). In addition, similar to 5FU, treatment with FUTP (**Fig. 5F**), as well as FUDR (**Supp. Fig. 5F**), also induced activation of the mTORC1 pathway and upregulation of ZNF598 and GIGYF2. This suggests that activation of mTOR pathway and upregulation of ZNF598 and GIGYF2 expression is a shared feature of 5FU-derived metabolites and independent of their abilities to incorporate into RNA and DNA, or to inhibit TS. Importantly, we also observed that, as was reported previously^46^, treatment with non-fluorinated UTP also rapidly increased mTORC1 activity and expression of ZNF598 and GIGYF2 (**Fig. 5G** & **Supp. Fig. 5G**). This indicates the presence of a shared mechanism of upregulation of mTOR activity among uridine derivatives, regardless of their fluorination status. We next examined if upregulation of ZNF598 and GIGYF2 is restricted to the conditions where mTOR is activated by uridine derivatives. We observed that chemical activation of mTOR with MHY1485^47^ in the absence of exogenous uridine-derived metabolites also increased ZNF598 and GIGYF2 expression (**Fig. 5H**).

Taken together, these data suggest the presence of an intrinsic mechanism of regulation of ZNF598 and GIGYF2 following mTOR activation that serves to avert the potential accumulation of collided ribosomes during mRNA translation activation. In the context of cancers such as CRC, wherein mRNA translation rate is elevated^48,49^, this intrinsic cellular mechanism could alleviate the cytotoxic repercussions of mTORC1 activation during acute 5FU treatment (**Fig. 6A**). Notably, using publicly available datasets for protein expression in normal and cancerous tissues^50^, we observed increased expression of ZNF598, GIGYF2, and ASCC3 proteins in CRC patient samples (**Fig. 6B-D**).

**Figure 6:**
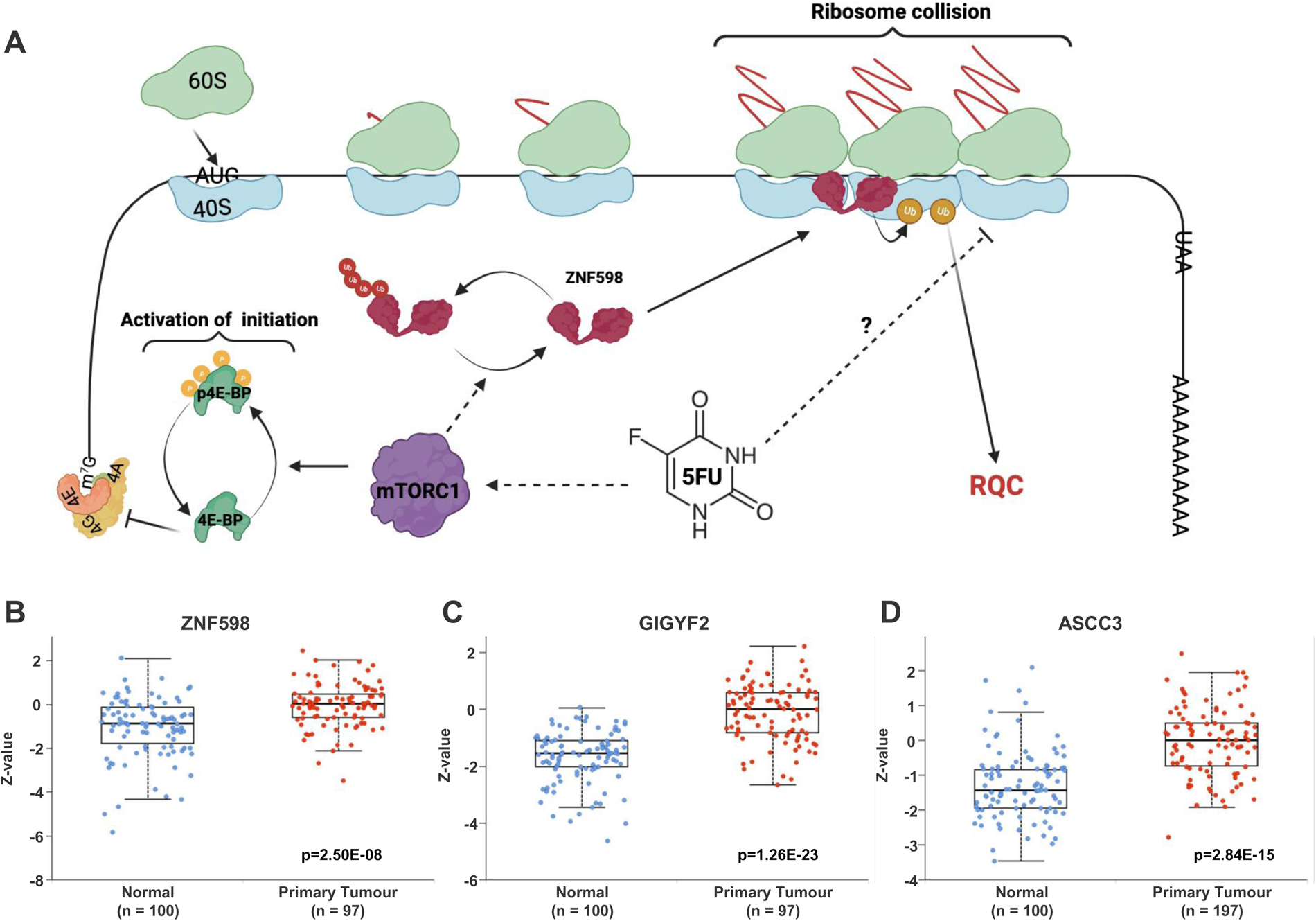
Enhanced mTOR activity couples elevated mRNA translation initiation with increased expression of RQC factors. (**A**) Proposed model for mechanism of 5FU-induced enhanced ribosome collisions and modulation of RQC activity. 5FU rapidly enhances mTOR activity by an unknown mechanism, leading to increased translation initiation via phosphorylation of factors such as 4E-BPs. Increased translation initiation elevates the risk of ribosome collisions. Fluorinated Uridines may also increase the risk of ribosome stalling through direct incorporation into the mRNA ORF or non-coding RNAs such as rRNAs, thereby further increasing the chances of ribosome collision. In parallel, activation of mTOR by 5FU also leads to stabilisation of ZNF598 and GIGYF2, which further bolsters RQC mechanism and contributes to mitigating the cytotoxic consequences of 5FU-induced ribosome stalling and collisions. (**B-D**) Protein expression levels of (**B**) ZNF598, (**C**) GIGYF2, and (**D**) ASCC3 in normal colon and colon adenocarcinoma samples. Data are extracted from The National Cancer Institute’s Clinical Proteomic Tumor Analysis Consortium (CPTAC)^50^ using UALCAN data analysis platform^77^. Z-values (number of standard deviations away from the mean) as well as p values are annotated for each protein.

## Discussion

Driven by the misconception that the transient nature of RNAs renders damage to them therapeutically inconsequential, the potential role of mechanisms such as RQC in the response of cancer cells to chemotherapy has been largely overlooked. Herein, we described a hitherto unknown mechanism of RNA-dependent 5FU toxicity, a drug which as noted above, incorporates into RNAs considerably more efficiently than it does into DNA.

Our study reveals a critical role for the mRNA translation machinery, mTOR signalling pathway, and RQC, a homeostatic mechanism of monitoring mRNAs to ensure translation integrity, in shaping the cellular response to 5FU. Challenging conventional beliefs, our data show that acute exposure to 5FU activates the mRNA translation machinery. This effect, previously obscured due to a tendency of the field to focus on the outcomes of 5FU cytotoxicity in long-term treatments, underscores the nuanced and dynamic influence of 5FU on mRNA translation. Importantly, we also discovered that 5FU treatment elevated the rate of ribosome collisions. Furthermore, we demonstrate that 5FU and its metabolites, as well as the non-fluorinated UTP, activate the mTOR pathway, which orchestrates the upregulation of key RQC factors ZNF598 and GIGYF2 along with the elevated rate of translation initiation and ribosome collisions. The RQC mechanism, activated in response to 5FU, efficiently resolves collided ribosomes and acts as a protective mechanism that abrogates the long-term 5FU-induced toxicity. Conversely, a defective RQC leads to accumulation of 5FU-induced collided ribosomes and enhanced 5FU-induced cell death. Together, our data highlight the potential for development of innovative synergistic treatments that simultaneously target the cellular RQC capacity and thereby augment the efficacy of 5FU-based standard of care anti-cancer therapy.

We provide robust evidence that acute 5FU treatment causes ribosome collisions. However, the exact mechanism by which 5FU induces ribosome stalling is still not clear. Our data revealed that through dysregulation of key signalling pathways, namely activation of mTORC1, acute 5FU treatment leads to enhanced mRNA translation initiation. This can pose an increased risk of ribosome collisions by elevating the ribosomal traffic on faulty mRNAs (e.g. aberrantly spliced), combined with aberrant translation events (e.g. stop codon readthrough), which are induced by 5FU^2,4,8^ and could potentially lead to ribosome stalling^29^. Furthermore, it is plausible that direct incorporation of 5FU metabolite FUTP in the open-reading frame (ORF) of the mRNAs or 5FU-induced changes in nucleotide modifications within the ORF could also impede ribosomes and result in stalling. While the ability of fluorinated uridines to directly impede ribosomes is not known, other types of modified uridine within the ORF including pseudouridine^16^ and 5-methyluridine^51^ were shown to stall ribosomes and triggers RQC^17^. Incorporation of 5FU-derived metabolites into non-coding RNAs, such as ribosomal RNAs (rRNAs) or transfer RNAs (tRNAs) may also contribute to the 5FU-induced ribosome stalling. Recent studies demonstrated the significant effects of incorporation of 5FU-derived metabolites into rRNAs on ribosome biogenesis and function^5,8^. In parallel, 5FU also alters tRNA modifications, stability, and function^7,52,53^, events which have, in other contexts, been linked to ribosome frameshifting and stalling^54,55^. While further analyses are required to deduce the exact mechanism by which 5FU induces ribosome stalling, our data clearly demonstrates that the enhanced rate of translation initiation induced by 5FU further exacerbates the rate of ribosome collisions. Consistent with this notion, blocking translation initiation with 4EGI-1 or Torin-1 markedly decreased the rate of ribosome collisions, presumably by lowering the ribosomal traffic and reducing the risk of collisions with stalled ribosome. Previous studies also demonstrated that blocking mTORC1-mediated translation attenuates the *Pelo* knockout RQC-deficiency phenotypes in mouse epidermal stem cells^31^.

RQC is a translation-dependent mechanism of quality control and more actively translated mRNAs are more prone to ribosome collisions and triggering RQC^20–22^, due to the higher chance of presence of trailing ribosomes that collide into a stalled ribosome. Metabolic stress and oxidative agents such as nitric oxide and reactive oxygen species, which are common in cancer cells and directly damage ribonucleotide bases, impede the transition of translating ribosomes and cause ribosome stalling^14,15,56,57^. Furthermore, following oncogenic transformation, overall translation activity is often elevated, partly due to mutation and activation of signalling pathways such as PI3K/mTOR^58^, which stimulate mRNA translation^12^. Consequently, cancers such as CRC that are reliant on elevated mRNA translation for tumour growth^48,49^ may exhibit dependency on intact RQC to mitigate the detrimental impacts of ribosome collisions. Our data demonstrate the synergistic enhancement of 5FU toxicity upon depletion of ZNF598 in two cell lines derived from two cancers treated with 5FU-based chemotherapy (colorectal cancer and pancreatic ductal adenocarcinoma). This provides a promising opportunity for enhancement of the 5FU anticancer efficacy by targeting the RQC machinery. However, further studies are required to fully characterise the impacts of genetics or pharmacological inhibition of various RQC factors on the response of cancer as well as non-cancer cells to 5FU based standard of care treatments. Importantly, our data also indicates that repression of mTOR activity and mRNA translation may render RQC-deficient cells less susceptible to the 5FU-derived ribosome collisions and cell death. While the precise impact of mTOR inhibition on 5FU efficacy is disputed^5,59,60^, our findings provides important insight into the potential cancer subtypes (i.e. high mTOR activity and mRNA translation) that may further benefit from a putative treatment that combines 5FU with RQC inhibition.

Significant progress has been made in our understanding of the mechanism of RQC^26^, its important role in maintenance of homeostasis^31,32^ and triggering global reprogramming of gene expression in response to stress^15,25,32,56,61^. Yet, important gaps exist in our understanding of how the RQC mechanism itself is regulated in response to environmental stimuli that impact cellular metabolism and mRNA translation, including the antimetabolites such as 5FU. Our data indicate the presence of a hitherto unknown cellular mechanism that increases expression of key RQC factors ZNF598 and GIGYF2 upon 5FU treatment and activation of mTOR signalling pathway. While the exact mechanism by which mTOR regulates the expression of ZNF598 and GIGYF2 is not completely understood, our data indicate a post-translational mechanism of regulation of protein stability (**Fig. 5**). Notably, a recent study suggested that mitochondrial stress, which is triggered upon 5FU treatment *in vitro* and *in vivo*^62–64^, leads to increased ZNF598 stability in a K63-linked ubiquitination dependent manner^65^. However, it is not clear how mitochondrial stress and/or mTOR activation could lead to ZNF598 or GIGYF2 deubiquitination and stabilisation upon acute 5FU treatment. Potential mechanisms include deactivation of an E3 ligase (plausibly the E3 activity of ZNF598 itself) or activation of a deubiquitinating (DUB) enzyme that leads to deubiquitylation and stabilisation of these proteins. Interestingly, depletion of USP9X – a DUB which reportedly enhances RQC by deubiquitylation and stabilisation of ZNF598^66^ – has been linked to increased sensitivity of CRC cells to 5FU^67,68^.

The mTOR pathway functions as a metabolic rheostat that integrates cellular response to various intracellular and extracellular signals (e.g., nutrients)^69^. Our data clearly demonstrate a robust and rapid, albeit temporal, activation of mTOR pathway upon treatment with 5FU and its metabolites, as well as the non-fluorinated UTP. Although the important role of mTOR in sensing the intracellular levels of purines has been demonstrated^70^, the available information on the mechanisms of monitoring the cellular pool of pyrimidines is scant. Thus, the mechanism by which 5FU and UTP lead to the activation of mTOR pathway remains to be understood. Interestingly, previous studies also suggested that ribosome stalling^31,71^ as well as impaired ribosomal biogenesis^72^ also activate mTORC1 signalling and stimulate translation initiation. It is therefore plausible that the impact of 5FU on ribosome biogenesis^5^ and the induction of ribosome stalling, as discussed above, may at least partially explain the rapid activation of mTOR signalling pathway by 5FU.

mTOR kinase forms two distinct protein complexes, mTORC1 and mTORC2, the former of which plays a pivotal role in regulation of mRNA translation. This is achieved via phosphorylation and regulation of activity of a group of translation factors and RNA-binding proteins, including 4E-BPs, S6Ks, and LARP1^12^. Active mTORC1 particularly favours translation of the highly abundant and efficiently translated nuclear encoded mitochondrial proteins^73^ and terminal oligopyrimidine (TOP) mRNAs^74,75^. As noted above, actively translated mRNAs are more prone to ribosome collisions and triggering RQC^20–22^. We propose that the swift stabilisation of ZNF598 and GIGYF2 proteins following mTOR activation serves as a mechanism to avert the potential accumulation of collided ribosomes during mRNA translation activation. In the context of cancers such as CRC wherein mTOR activity is frequently upregulated^76^, this intrinsic cellular mechanism likely aids cancer cells to alleviate the cytotoxic repercussions of mTORC1 activation during acute 5FU treatment. Our examination of publicly available datasets revealed heightened expression of ZNF598, GIGYF2, and ASCC3 in CRC in comparison to non-tumour tissues. Further investigations are essential to elucidate the impact of mTOR activation, such as mutations in components of the PI3K/mTOR pathway, on the increased expression of these factors and their role in tumour maintenance.

Altogether, we demonstrate that contrary to previous assumptions, acute 5FU treatment leads to activation of mRNA translation as well as induction of ribosome collisions. We show the critical role of the RQC mechanism in resolving the 5FU-induced ribosome collisions and mitigating their deleterious consequences, including a profound impact on overall cellular viability and clonogenic survival in response to this keystone chemotherapeutic. This report presents compelling evidence pointing to the existence of a previously unrecognized mechanism of coupling mTOR-mediated activation of mRNA translation to the increased expression of key RQC factors. This mechanism may hold broad significance across various biological and pathological contexts involving mTOR-mediated upregulation of mRNA translation, including development, differentiation, and tumorigenesis. Besides providing fundamental insights into the mechanisms of regulation of RQC activity, this work elucidates a mechanism of cellular response to the RNA-dependent toxicity of 5FU, the mainstay component of the chemotherapeutic treatments for several common and highly lethal cancers. Thus, our findings may lead to potentially significant improvement in utility of 5FU-based anti-cancer treatments by leveraging the cancer cell dependency on RQC.

## Materials & Methods

### Cell lines and culture conditions

HCT116 human colorectal cancer and SUIT-2 human pancreatic ductal adenocarcinoma cells were grown in McCoy’s 5A medium (Gibco, Cat. # 26600080) supplemented with 10% v/v dialyzed Foetal Bovine Serum (FBS) (Sigma, Cat. # F0392), 1 mM Sodium Pyruvate (Gibco, Cat. # 11360039), 100 U/ml penicillin, 100 µg/ml streptomycin (Gibco, Cat. # 15070063). HEK293 (Thermo Fisher Scientific) were cultured in DMEM (Dulbecco’s Modified Eagle Medium; Gibco, Cat. # 41965039) supplemented with 10% FBS (Gibco, Cat. # 10270106) and 100 U/ml penicillin and 100 µg/ml streptomycin (Gibco, Cat. # 15070063). All cells were maintained at 37℃, in a humidified atmosphere with 5% CO_2_ and regularly tested for presence of mycoplasma contamination using mycoplasma detection kit (abm, Cat. # G238).

### Immunofluorescence

HCT116 cells were seeded onto poly-L-lysine coated cover slips in 6 well plates. Cells were seeded at a density of 1.8 x 10^5^ and allowed to adhere for 24 h. Cells were then treated with 2.5 µM 5FU (Merck, Cat. # 343922) or vehicle for the indicated time points. After treatment, the cells were washed in PBS and fixed in pre-chilled 70% ethanol for 30 min at -20°C. For RNase digestion, fixed cells were incubated with 400 U of RNase I (Thermo Fisher Scientific, Cat. # EN0601) in PBS for 1 h at 37°C, followed by washes with PBS post-digestion. Prior to 5FU detection, depurination of the cells was performed with 2 M hydrochloric acid for 30 min at room temperature (RT) followed by 3 washing with PBS. The cells were then blocked with a bovine serum albumin/normal goat serum buffer containing 0.5% Triton X-100 for 1 h. Subsequently, cells were further blocked with a streptavidin/biotin blocking kit (Vector Laboratories, Cat. # SP2002). Next, 5FU uptake into cells was captured using an anti-BrdU antibody (BU-33; Merck) at 1:150 dilution in 1% BSA/PBS. After 1 h incubation at RT with the primary antibody, 5FU was detected using an anti-mouse biotinylated secondary antibody (Thermo Fisher Scientific, Cat. # A10519) at 1:200 dilution in 1% BSA/PBS for 1 h at RT followed by incubation with streptavidin-conjugated 488 (Thermo Fisher Scientific, Cat. # S11223) at 1:1000 dilution in PBS for 1 h at RT. Cells were then counterstained with DAPI, and coverslips mounted onto microscopy slides. Images were taken using confocal microscopy (Olympus Spinning Disk) at 40x magnification. 5FU fluorescence was analysed with ImageJ software (NIH).

### Generation of ZNF598-KO HCT116 and SUIT-2 cells

The pSpCas9(BB)-2A-GFP (Addgene, Cat. 48138) plasmid was linearized using BbsI (Thermo Fisher Scientific, Cat. # ER1011) and the oligodeoxynucleotides encoding guide RNA (5′-ACCGCTGCTCTACCAAGATG) for targeting the coding region of ZNF598 gene were ligated. The ligation reaction mix was transformed into *E. coli* Dh5∝ strain and after transformation, the guide sequence containing pSpCas9(BB)2A-GFP plasmids were isolated and verified by Sanger sequencing using the U6 promoter forward primer. To generate the knockout cells, 250,000 cells were plated into 12 well plates and transfected with the guide sequence containing pSpCas9(BB)-2A-GFP plasmid using Lipofectamine 3000 (Thermo Scientific, Cat. # 11668019). 24 h after transfection, GFP positive cells were sorted by FACS into 96-well plates and cultivated until colonies were obtained. Clonal cell lines were analyzed by WB for the absence of the protein. Clones showing lack of ZNF598 protein expression were verified by Sanger sequencing using ZNF598 specific primer pairs: GCAGCTCCTGAGCGGGGAG (forward) and CCGGACCTCAGTGAGGCAAAGT (reverse).

### Lentivirus shRNA packaging

Lentivirus pseudovirions were produced by transfecting HEK293T cells using Lipofectamine 2000 (Thermo Fisher Scientific, Cat. # 11668027) and 1.25 µg shRNA plasmid along with equivalent amount of psPAX2 (Addgene, plasmid 12260) and pMD2.G (Addgene, plasmid 12259) in 3:1 ratio in 6 well plates. 48 h post-transfection, media was collected from which pseudovirions were purified by filtration (Filtropur S, 0.2 µm, Sarstedt, Cat. # 83.1826.001) after a brief centrifugation (1500 x g, 5 min). The following shRNAs were used in this study: Non-Targeting shRNA Controls (Sigma, Cat. # SHC002), shZNF598#1 (Sigma, Cat. # TRCN0000073158), shZNF598#2 (Sigma, Cat. # TRCN0000073162), shZNF598#3 (Sigma, Cat. # TRCN0000222610).

### RNA extraction and quantitative RT-PCR

Total RNA was isolated using TRIzol reagent (Thermo Fisher Scientific, Cat. # 15596026) as per the manufacturer’s protocol. 1 µg purified total RNA was treated with DNase I (Thermo Fisher Scientific, Cat. # EN0521) prior to reverse transcription using SuperScript™ III Reverse Transcriptase (Thermo Fisher Scientific, Cat. # 18080085) and random hexamers. The following primers were used for PCR reactions: ZNF598-Forward: AAAGGTGTACGCATTGTACAGG, ZNF598-Reverse: CTCCAGGTCCCCGAAGAG, GIGYF2-Forward: TCTGTGGGTCAGGAATTTGG, GIGYF2-Reverse: GACATCTGACCACAACCAAAGA, 28S rRNA-Forward: CGATGTCGGCTCTTCCTATC, 28S rRNA-Reverse: TCCTCAGCCAAGCACATACA. Quantitative PCR was performed using LightCycler® 480 Instrument II (Roche, Cat. # 05015278001) using the LightCycler 480 SYBR Green I Master mix (Roche, Cat. # 04887352001), according to the manufacturer’s protocol.

### Western blotting

Cells were washed with pre-chilled PBS and detached from the plates using plastic scrapers. After centrifugation, cell pellets were lysed in RIPA buffer (50 mM Tris-HCL pH 7.4, 150 mM NaCl, 2 mM EDTA, 1% NP-40, 0.1% SDS) supplemented with 1 mM Sodium orthovanadate (Na_3_VO_4_) and protease inhibitors (Roche, Cat. # 11836170001). Total protein concentration was measured by Bradford protein assay (Bio-Rad, Cat. # 5000006) and 40 µg of total protein was mixed with 5x loading buffer (0.25% Bromophenol blue, 0.5 M DTT, 50% Glycerol, 10% SDS, and 0.25 M Tris-Cl pH 6.8). Samples were boiled for 3 min, followed by incubation on ice for 10 min and loaded on SDS-PAGE gel and wet transfer onto PVDF membrane (Merck, Cat. # IPFL00010). Thereafter, membranes were blocked with 5% BSA (Thermo Fisher Scientific, Cat. # A9647) at room temperature for 1 h and incubated with the indicated primary antibodies overnight at 4°C and secondary antibody for 2 h at room temperature. All blots were scanned, and images were captured using the Odyssey system (LI-COR, Cat. # ODY-1540). When necessary, brightness and contrast adjustments were applied evenly across the entire image. Uncropped versions of all western blot images are presented in **Supp. Fig. 6-18**. For quantification of the total or phospho-proteins, the intensity of each band was measured with ImageJ software (NIH) and normalised against the total protein (in case of phospho-proteins) or the loading control (β-actin) after subtracting immunoblot background intensity.

The following antibodies were used in this study: rabbit anti-phospho-RPS6 (S240/244) (Cell Signalling, Cat. # 5364), rabbit anti-RPS6 (Cell Signalling, Cat. # 2217), rabbit anti-phospho-4E-BP1 (Cell Signalling, Cat. # 2855), rabbit anti-4E-BP1 (Cell Signalling, Cat. # 9644), rabbit anti-phospho-eEF2 (Cell Signalling, Cat. # 2331), rabbit anti-eEF2 (Cell Signalling, Cat. # 2332), rabbit anti-phospho-eIF2α (Abcam, Cat. # ab32157), rabbit anti-eIF2α (Cell Signalling, Cat. # 5324), rabbit anti-phospho-p38 MAPK (Cell Signalling, Cat. # 9211), rabbit anti-p38 MAPK (Cell Signalling, Cat. # 9212), rabbit anti-phospho-eIF4E (Abcam, Cat. # ab76256), mouse anti-eIF4E (BD Biosciences, Cat. # 610270), rabbit anti-eS10 (Abcam, Cat. # ab151550), rabbit anti-RACK1 (Cell Signalling, Cat. # 4716), rabbit anti-ZNF598 (Invitrogen, Cat. # 703601), rabbit anti-GIGYF2 (Proteintech, Cat. # 24790-1-AP), mouse anti-puromycin (Merck, Cat. # MABE343), rabbit anti-PARP (Cell Signalling, Cat. # 9542), rabbit anti-EDF1 (Abcam, Cat. # ab174651), rabbit anti-ASCC3 (Proteintech, Cat. # 17627-1-AP) mouse anti-β-Tubulin (Sigma, Cat. # T4026), rabbit anti-GAPDH (Proteintech, Cat. # 10494-1-AP), mouse anti-β-actin (Sigma, Cat. # A5441), rabbit anti-Vinculin (Cell Signalling, Cat. # 13901S), IRDye® 680RD Donkey anti-Mouse IgG (LI-COR, Cat. # 926-68072), and IRDye® 800CW Donkey anti-Rabbit IgG (LI-COR, Cat. # 926-32213).

### Polysome profiling

Cells were maintained at maximum 60-70% confluency. After pre-treatment with Cycloheximide (100 μg/ml; Sigma, Cat. # 01810) for 5 min, cells were collected by centrifugation at 4°C for 5 min and lysed in 500 μl hypotonic buffer containing 5 mM Tris-HCl, pH 7.5, 2.5 mM MgCl2, 1.5 mM KCl, complete EDTA-free protease inhibitor cocktail (Roche, Cat. # 04693159001), 100 μg/ml Cycloheximide, 2 mM DTT, 200 U/ ml RiboLock RNase Inhibitor (Thermo Fisher Scientific, Cat # EO0382), 0.5% v/w Triton X-100, and 0.5% v/w sodium deoxycholate. The lysates were cleared by centrifugation at 20,000 × g for 5 min at 4°C. 400 µg of RNA, measured by NanoDrop 2000 (Thermo Fisher Scientific), were loaded onto 10%–50% sucrose gradients. The samples were sedimented by velocity centrifugation at 36,000 x rpm for 2 h at 4°C using SW40Ti rotor in Optima L-80XP ultracentrifuge (Beckman Coulter). Absorbance at 254 nm was measured from lower to higher sucrose gradients using an ISCO gradient fractionation system and the optical density at 254 nm was continuously recorded with a Foxy JR Fractionator (Teledyne ISCO).

### Detection of ribosome collisions by micrococcal nuclease treatment

HCT116 cells were treated with 5FU for the indicated time-points. Cells were pre-treated with Cycloheximide (100 μg/ml) for 5 min before harvesting at maximum confluency of 60-70%, collected by centrifugation at 4°C for 5 min and lysed in 500 μl hypotonic buffer containing 5 mM Tris-HCl, pH 7.5, 2.5 mM MgCl_2_, 1.5 mM KCl, complete EDTA-free protease inhibitor cocktail, 100 μg/ml Cycloheximide, 2 mM DTT, 200 U/ ml RiboLock RNase Inhibitor, 0.5% v/w Triton X-100, and 0.5% v/w sodium deoxycholate. Indicated cells were pre-treated with 15 nM Torin-1 (Sigma, Cat. # 475991) or 25 μM 4EGI-1 (Selleck Chemicals, Cat. # S7369) for 24 h before incubation with 2.5 μM of 5FU for the indicated time-points. The lysates were cleared by centrifugation at 20,000 × g for 5 min at 4°C. CaCl2 was added to the lysate to a final concentration of 1 mM, and 500 µg of the lysate was digested with 1000 U micrococcal nuclease (NEB, Cat. # M0247S) for 30 min at 22°C. Digestion by micrococcal nuclease was stopped by adding 2 mM EGTA. Afterwards, RNA-equivalent amounts of lysate (were resolved on 10-50% sucrose gradients and absorbance at 254 nm was measured from lower to higher sucrose gradients.

### Cell growth assay measurement

HCT116 or SUIT-2 cells were seeded at 25,000 cells per well in 6-well plates in complete medium for the indicated times. Cells were trypsinized and stained with Trypan Blue for 2 min and counted under the microscope using Haemocytometer. The process was repeated for 5 days to monitor the comparative cell growth.

### Colony formation assay

HCT116 and SUIT-2 cells were seeded at 400 cells per well in 6-well plates. To study differential colony formation after drug treatment, cells were treated with respective doses of 5FU and maintained at 37°C and 5% CO_2_. After 7 days, fresh media containing respective doses of 5FU was replenished. After 12 days, cells were rinsed briefly with PBS before staining with colony fixation-staining solution (crystal violet 0.4% w/v in 95% Methanol) for 5 minutes, followed by two brief washes in ice cold water and left to dry overnight. Afterwards, the number of colonies (diameters >200 µm) was counted using the Oxford Optronix GelCount™ system (v1.1.2.0; Oxford Optronix).

### Dose response assays and IC50 measurement

HCT116 and SUIT-2 cells were seeded at 2,000 cells per well in 96-well plates in complete media containing dialyzed FBS. The following day fresh media containing serial dilutions of 5FU (Merck, Cat. # 343922), FUDR (Sigma, Cat. # F0503), FUTP (Biorbyt, Cat. # orb64970) or Gemcitabine (Sigma, Cat. # G6423) were added. 72 h later cell viability was measured using CellTiter-Glo® Luminescent Cell Viability Assay reagent (Promega, Cat. # G7572) according to the manufacturer’s protocol. The comparative luminescence was measured on the Synergy Microplate reader (Biotek, Synergy HTX).

### SUrface SEnsing of Translation (SUnSET) assay

HCT116 and SUIT-2 cells were seeded at 2.5 x 10^5^ cells per well in 6-well plates and treated with 5FU (2.5 μM) for indicated times. Puromycin dihydrochloride (Sigma, Cat. # P8833) was added to the media at 2 μg/ml final concentration for 20 min prior to cell lysis with RIPA buffer. A group pre-treated with 100 μg/ml of Cycloheximide for 5 min was used as an additional negative control. 10 µg of cell lysates were loaded in a 10% SDS-PAGE gel, followed by western blotting, and probing with anti-Puromycin antibody. The total signal density was quantified using Image J software (NIH).

### Pulse labelling with the methionine analogue L-homopropargylglycine

Cells were seeded at 1,000 cells per well in 96-well plates and treated with 2.5 μM of 5FU for the indicated time, followed by incubation in methionine-free medium containing the methionine analogue L-homopropargylglycine (HPG) for 30 min. Cells were next fixed with 4% paraformaldehyde for 15 min and permeabilized using 0.5% Triton X-100 solution. The differential level of nascent protein synthesis was quantified using Click-iT™ HPG Alexa Fluor™ 488 Protein Synthesis Assay Kit (Invitrogen, Cat. # C10428) according to the manufacturer’s protocol on a BMG FLUOstar Omega Microplate Reader. The signal was normalized against the unstained samples for each set of experiments.

### Statistical Analyses

Statistical analyses were performed using Prism 6 (GraphPad). Error bars represent standard deviation (SD) from the mean of at least 3 independent replicates and individual data points are depicted in bar graphs. Statistical significance was set a priori at 0.05. Only one observation per sample was collected.

## Acknowledgment

We thank Yumna Azam, Purnima Kovuri, Aishwarya Khadanga, and Lilas Alboushi for technical assistance and Dr. Emma Kerr for discussions. This work was supported by an Academy of Medical Sciences Springboard Award (SBF007\100026), The Royal Society Research Grant (RGS\R1\221075), and Patrick Johnston Research Fellowship to S.M.J. and The Royal Society Research Grant (RGS\R2\222149) and the Division of Cancer Sciences, University of Manchester to J.R.P.K. T.M. is supported by a PhD studentship from Brainwaves Northern Ireland.

## Authors’ contribution

Conceptualization: S.C. and S.M.J.; Investigation & Methodology: S.C, P.N., N.S., A.G., K.C., A.H., P.H. S, T.N.S., T.M., K.D., and Z.V.S.; Reagent contribution: A.G., T.T., S.M.D., D.B.L.; Writing original draft, S.C., S.M.J; Review & Editing, all authors; Supervision, S.M.J.

---

**Supplementary Figure 1:**
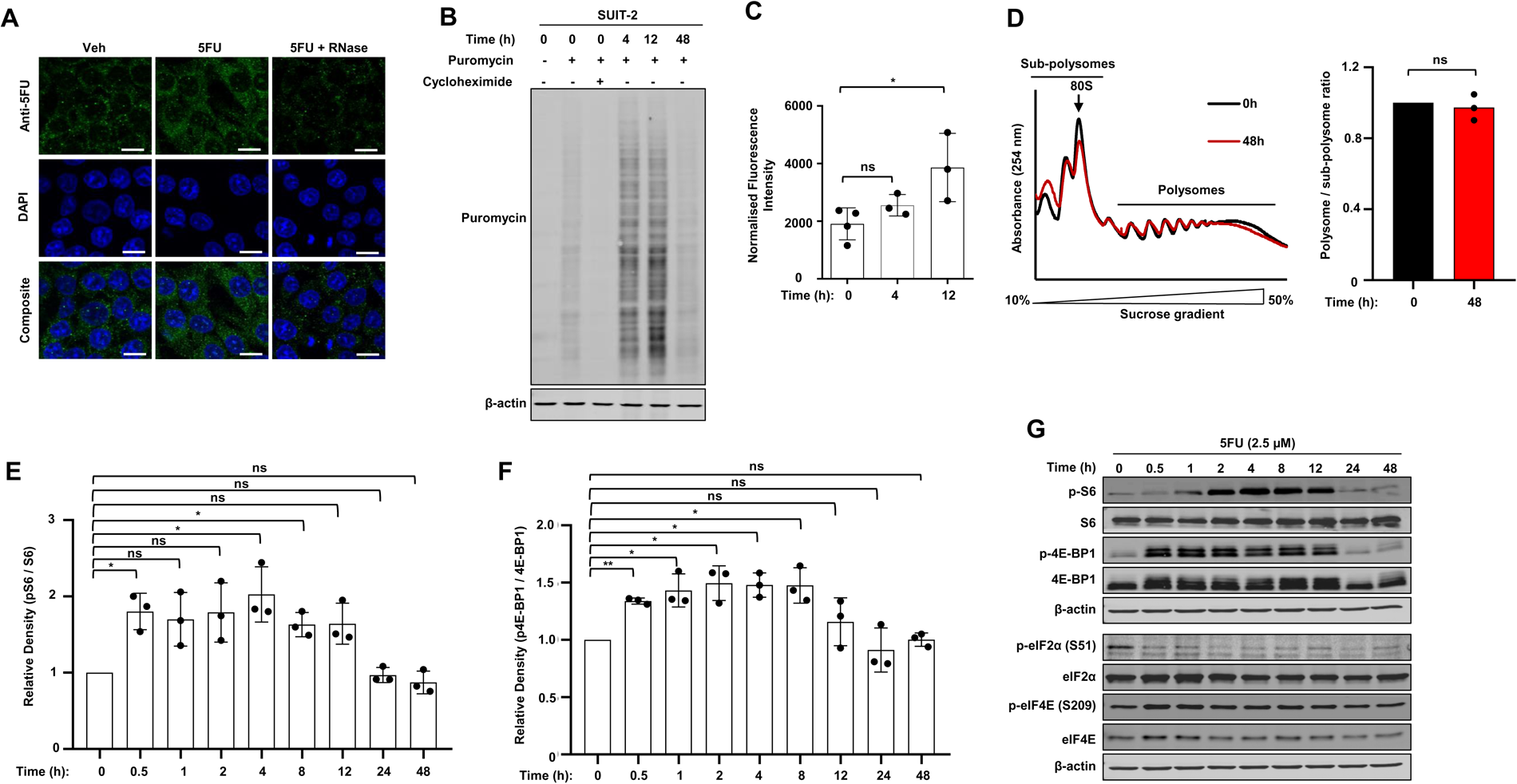
Acute 5FU treatment enhances mRNA translation. Related to Figure 1. (**A**) HCT116 cells were seeded onto coverslips and allowed to adhere for 24 h for immunofluorescence analysis of 5FU incorporation. Cells were treated with 5FU (2.5 µM) for 24 h timepoints. Cells in the Ribonuclease treated group were incubated with 400 U of RNase for 1 h, followed by washes with PBS and stained for 5FU (green) using anti-BrdU antibody and counterstained in the nucleic acid dye DAPI (blue). Scale bar = 45 µm. (**B**) Assessment of nascent protein synthesis by Surface Sensing of Translation (SUnSET) assay in SUIT-2 pancreatic ductal adenocarcinoma cells treated with 5FU (2.5 µM) for the indicated times. (**C**) Quantification of new protein synthesis by pulse-labeling with the methionine analogue L-homopropargylglycine (HPG). SUIT-2 cells were treated with 5FU (2.5 µM) for the indicated times followed by incubation in methionine-free medium containing the HPG for 30 min and measurement of nascent protein synthesis with Click-iT™ HPG Alexa Fluor™ 488 Protein Synthesis Assay Kit. The signal was normalized against the unstained samples for each group. Data are presented as mean ± SD; n=3 independent replicates; *p< 0.05, two-tailed student’s t-test. (**D**) *Left.* Analysis of general mRNA-ribosome association in HCT116 cells treated with 5FU (2.5 µM) for 48 h using polysome profiling assay. *Right.* Representative quantification of the polysome/sub-polysome ratio in control and 5FU treated cells. Data are presented as mean ± SD; n=3 independent replicates; two-tailed student’s t-test. (**E & F**) Quantification of expression of phopsho-RPS6 (pS6) and p4E-BP1 in Fig. 1E. Bar graphs represents the changes in (**E**) pS6 normalised to total RPS6 and (**F**) p4E-BP1 normalised to total 4E-BP1 relative to the 0 h control. Densitometric quantification was performed by ImageJ. Data are presented as mean ± SD; n=3 independent replicates; ns=non-significant, *p<0.05, **p<0.01, two-tailed student’s t-test. (**G**) Western blot analysis of effects of treatment with 5FU (2.5 µM) for the indicated times on main signalling pathways regulating mRNA translation machinery in SUIT-2 cells.

**Supplementary Figure 2:**
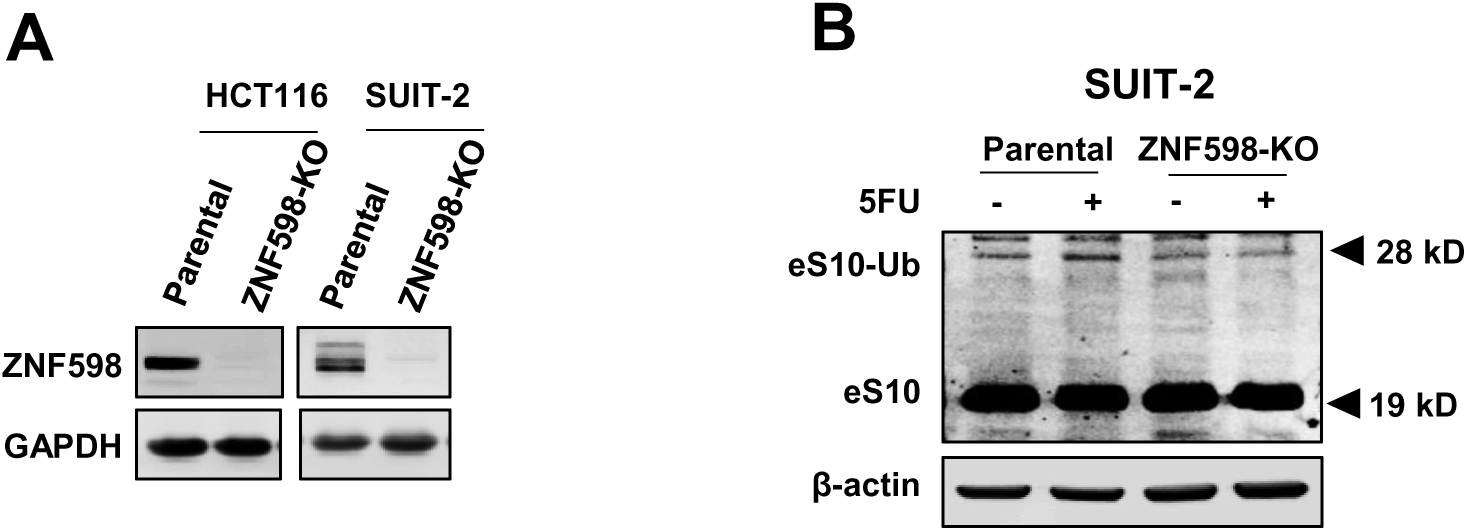
ZNF598 depletion results in dysregulated response to 5FU-treatment. Related to Figure 2. (**A**) Western blot analysis of ZNF598 expression in parental (WT) and CRISPR-Cas9 mediated ZNF598-KO HCT116 and SUIT-2 cells. GAPDH was used as a loading controls. (**B**) Western blot analysis of mono-ubiquitination of eS10 upon 8 h 5FU treatment (2.5 µM) in parental and ZNF598-KO SUIT-2 cells. β-actin was used as a loading controls.

**Supplementary Figure 3:**
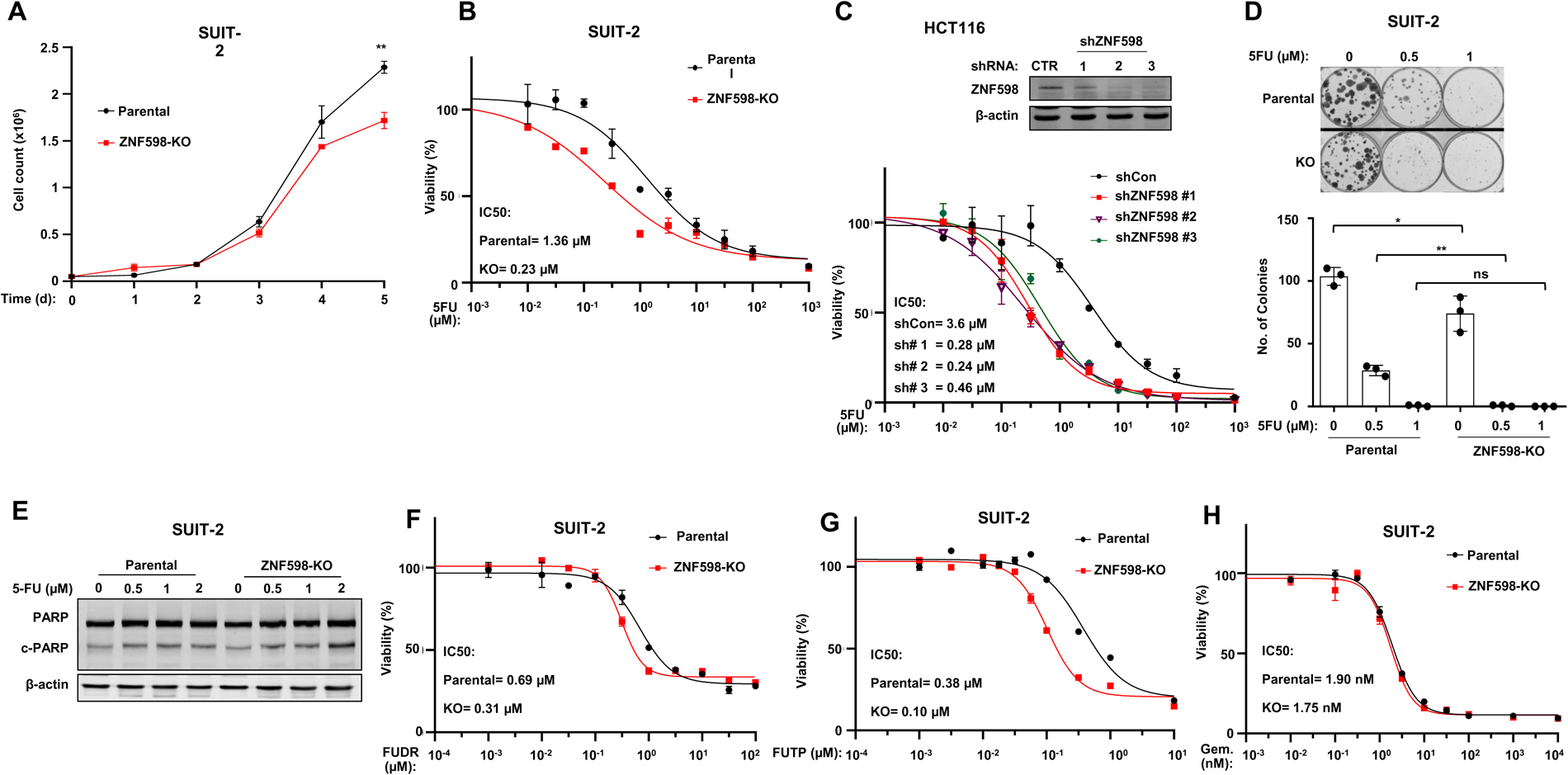
ZNF598-KO renders cancer cells more sensitive to 5FU induced cell death. Related to Figure 3. (**A**) Cell growth assay with parental and ZNF598-KO SUIT-2 cells. Cells were harvested after the indicated time points and cell numbers determined using a haemocytometer. Data are shown as mean ± SD; n = 3 independent replicates; **p<0.01, two-tailed student’s t-test. (**B**) Dose-response assay for measurement of sensitivity of parental and ZNF598-KO SUIT-2 cells to 5FU. 72 h post-treatment cell viability was measured using CellTiter-Glo® Luminescent Cell Viability Assay. (**C**) *Top:* Western blot analysis of ZNF598 knockdown with 3 different shRNAs (shZNF598# 1-3) in HCT116 cells. β-actin was used as loading control. *Bottom:* Dose-response assay for measurement of sensitivity of control (shCTR) and ZNF598 Knockdown (shZNF598# 1-3) HCT116 cells to 5FU. 72 h post-treatment cell viability was measured using CellTiter-Glo® Luminescent Cell Viability Assay. (**D**) *Top*: Colony formation assay with parental and ZNF598-KO SUIT-2 cells 12 days post-seeding. Cells were treated with the indicated concentration of 5FU. *Bottom*: Quantification of changes in the number of colonies. Colonies with a diameter >200 µm were counted. Data are presented as mean ± SD; n=3 independent replicates; *p<0.05; **p<0.01, two-tailed student t-test. (**E**) Western blot analysis of expression of cleaved PARP (c-PARP) using lysates derived from parental and ZNF598-KO SUIT-2 cells treated with the indicated doses of 5FU for 72 h. β-actin was used as loading control. (**F-H**) Dose-response assay for measurement of sensitivity of parental and ZNF598-KO SUIT-2 cells to (**F**) FUDR, the precursor of the DNA-incorporating 5FU metabolite, (**G**) the RNA-incorporating 5FU metabolite FUTP, and (**H**) Gemcitabine. 72 h post-treatment cell viability was measured using CellTiter-Glo® Luminescent Cell Viability Assay.

**Supplementary Figure 4:**
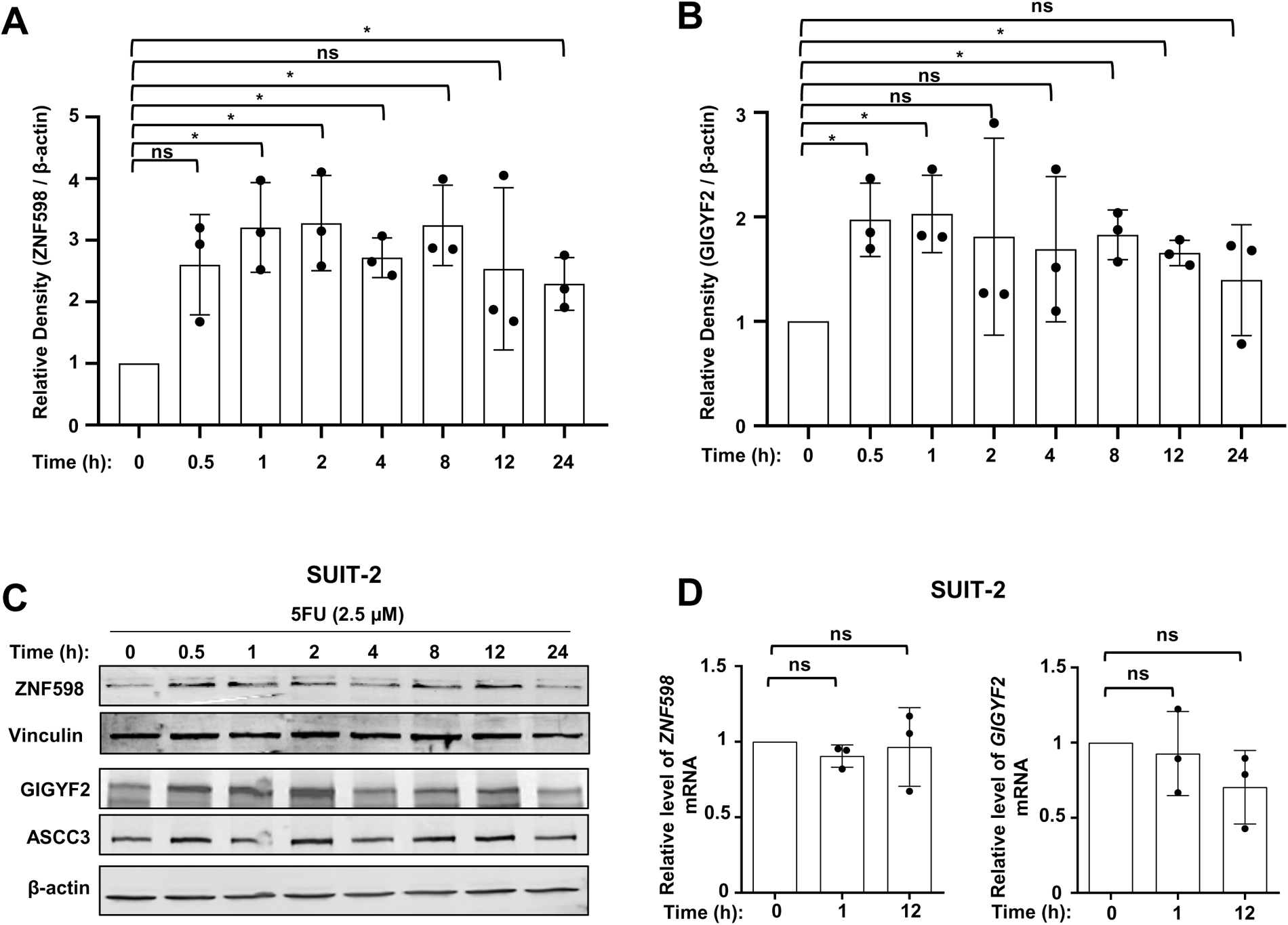
Upregulation of ZNF598 and GIGYF2 protein expression in response to 5FU treatment. Related to Figure 5; (**A-B**) Quantification of expression of (**A**) ZNF598 and (**B**) GIGYF2 proteins in Fig. 5A. Bar graphs represents the changes in ZNF598 and GIGYF2 normalised to β-actin relative to the 0 h control. Densitometric quantification was performed by ImageJ. Data are presented as mean ± SD; n=3 independent replicates; ns=non-significant, *p<0.05, two-tailed student’s t-test. (**C**) Western blot analysis of expression of indicated proteins in lysates derived from SUIT-2 cells treated with 5FU (2.5 µM) for the indicated time points. (**D**) Quantitative RT-PCR analysis of expression of *ZNF598* and *GIGYF2* mRNAs in SUIT-2 cells treated with 5FU (2.5 µM) for the indicated times. Data are presented as mean ± SD; n=3 independent replicates; ns=non-significant; two-tailed student’s t-test.

**Supplementary Figure 5:**
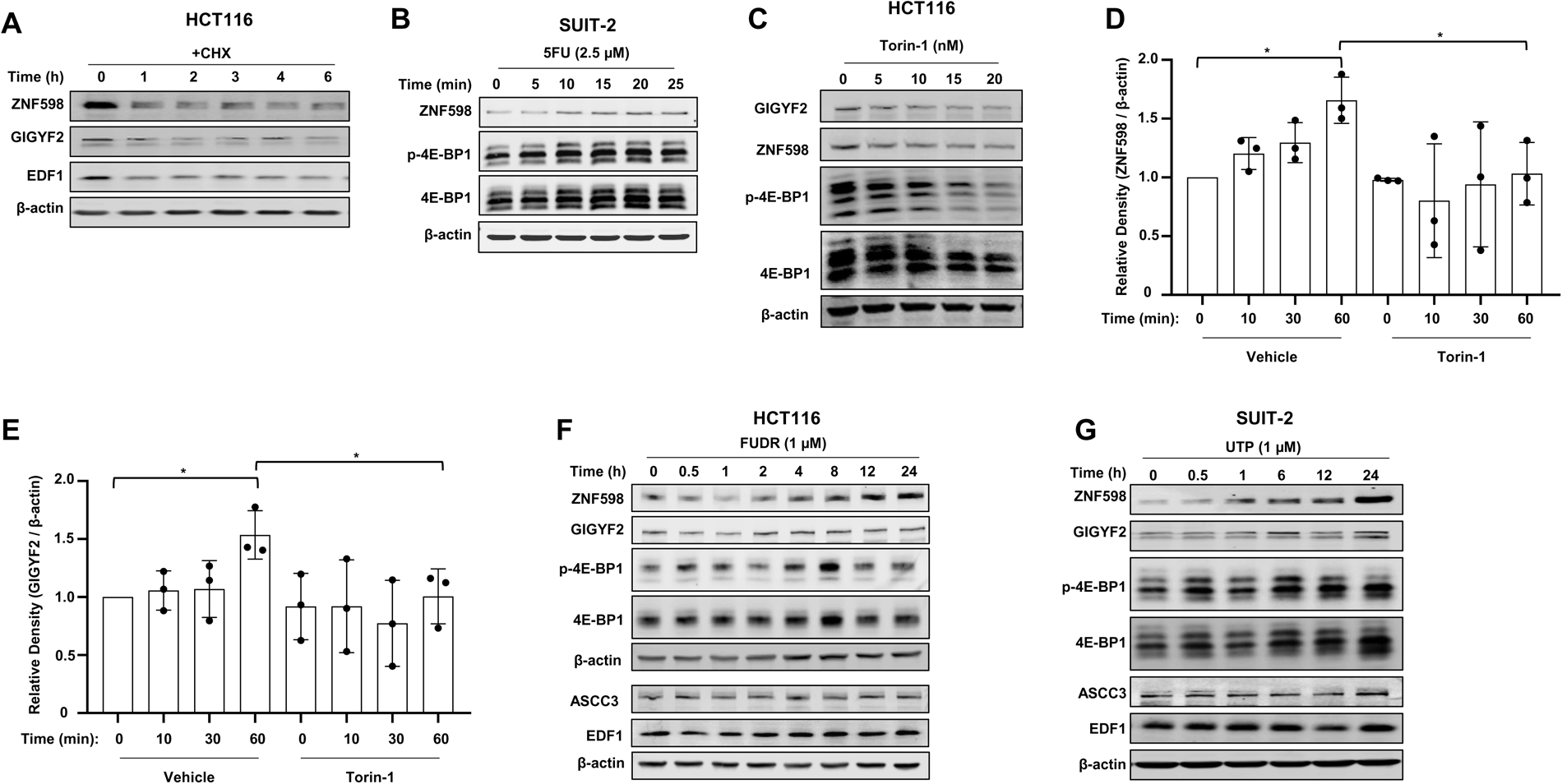
5FU-derived metabolites and Uridine induce ZNF598 and GIGYF2 expression. Related to Figure 5. (**A**) Western blot analysis of expression of the indicated proteins in lysates derived from HCT116 cells treated with 100 µg/ml Cycloheximide (CHX) for the indicated time points. (**B**) Western blot analysis of expression of indicated proteins in lysates derived from SUIT-2 cells treated with 5FU (2.5 µM) for the indicated time points. (**C**) Western blot analysis of expression of indicated proteins in lysates derived from HCT116 cells treated with the indicated doses of Torin-1 for 24 h. (**D-E**) Quantification of expression of (**D**) ZNF598 and (**E**) GIGYF2 proteins in Fig. 5E. Bar graphs represents the changes in ZNF598 and GIGYF2 normalised to β-actin relative to the 0 h control in each treatment group (Vehicle or Torin-1). Densitometric quantification was performed by ImageJ. Data are presented as mean ± SD; n=3 independent replicates; ns=non-significant, *p<0.05, two-tailed student’s t-test. (**F**) Western blot analysis of lysates derived from HCT116 cells treated with FUDR (1 µM) for the indicated time points. To avoid excessive non-specific signals due to repeated re-probing, identical samples were run on 2 separate gels. (**G**) Western blot analysis of expression of the indicated proteins in lysates derived from SUIT-2 cells treated with UTP (1 µM) for the indicated time points.

**Supplementary Figure 6.**
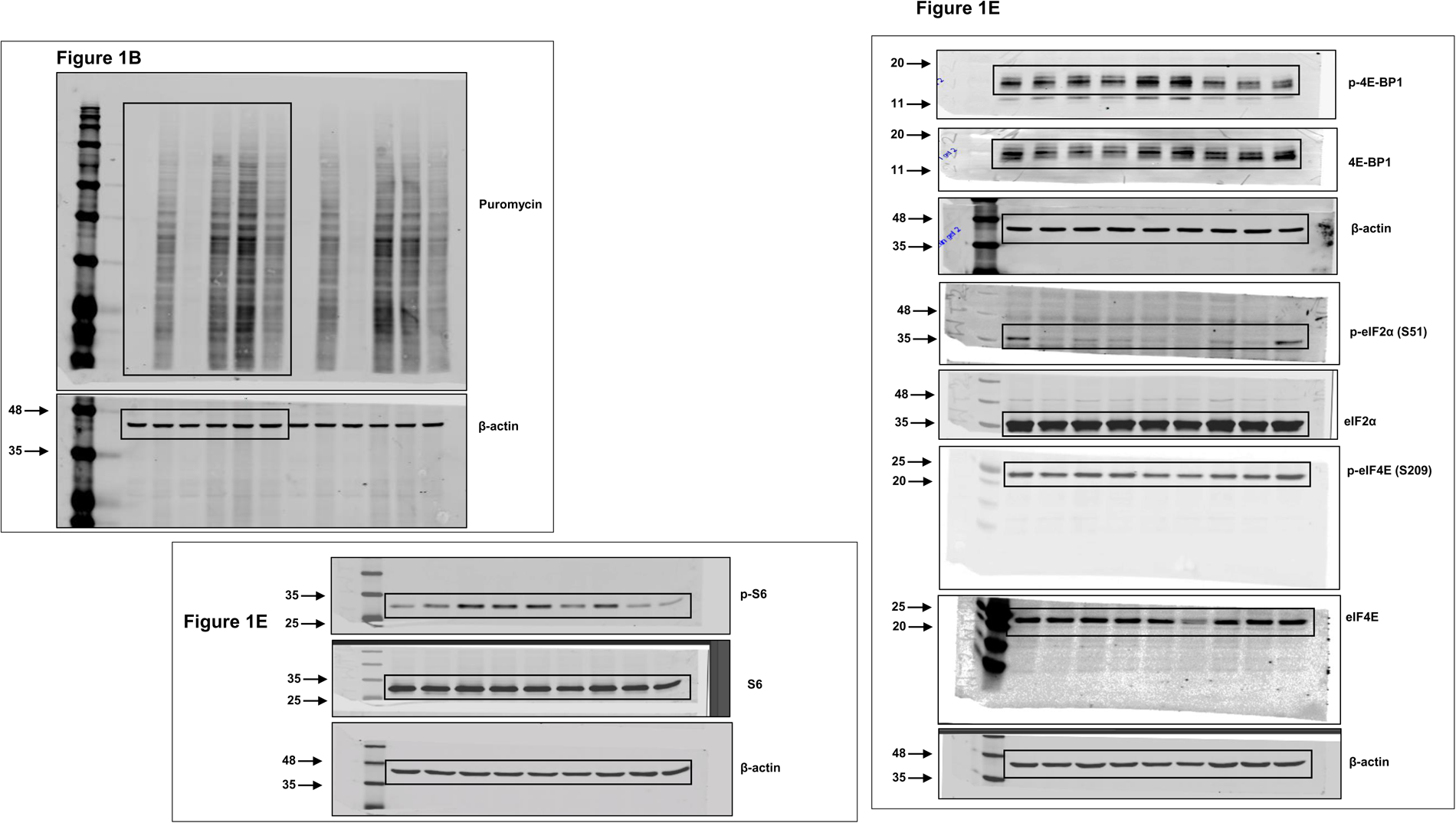
Uncropped images of blots used in Figure 1.

**Supplementary Figure 7.**
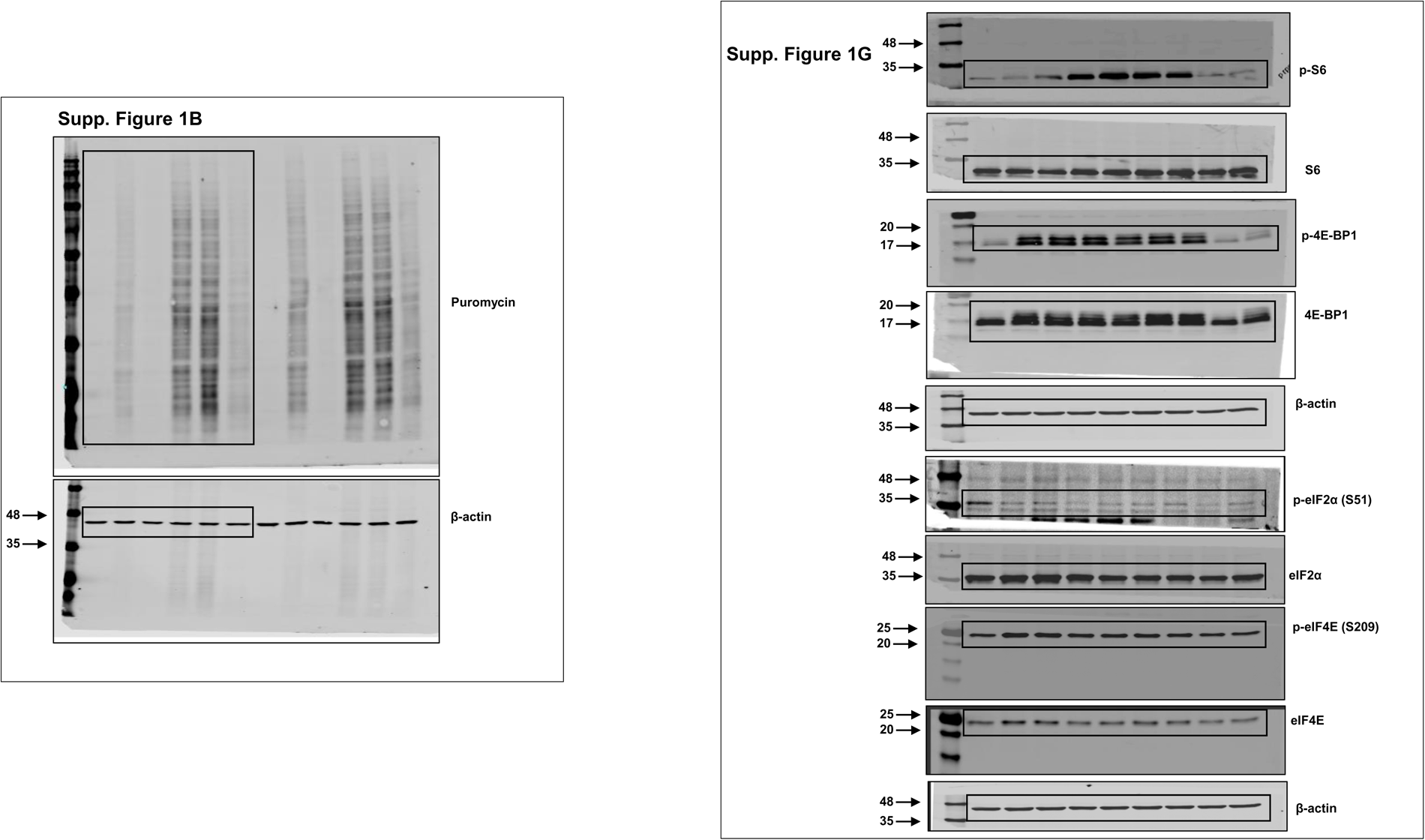
Uncropped images of blots used in Supp. Figure 1.

**Supplementary Figure 8.**
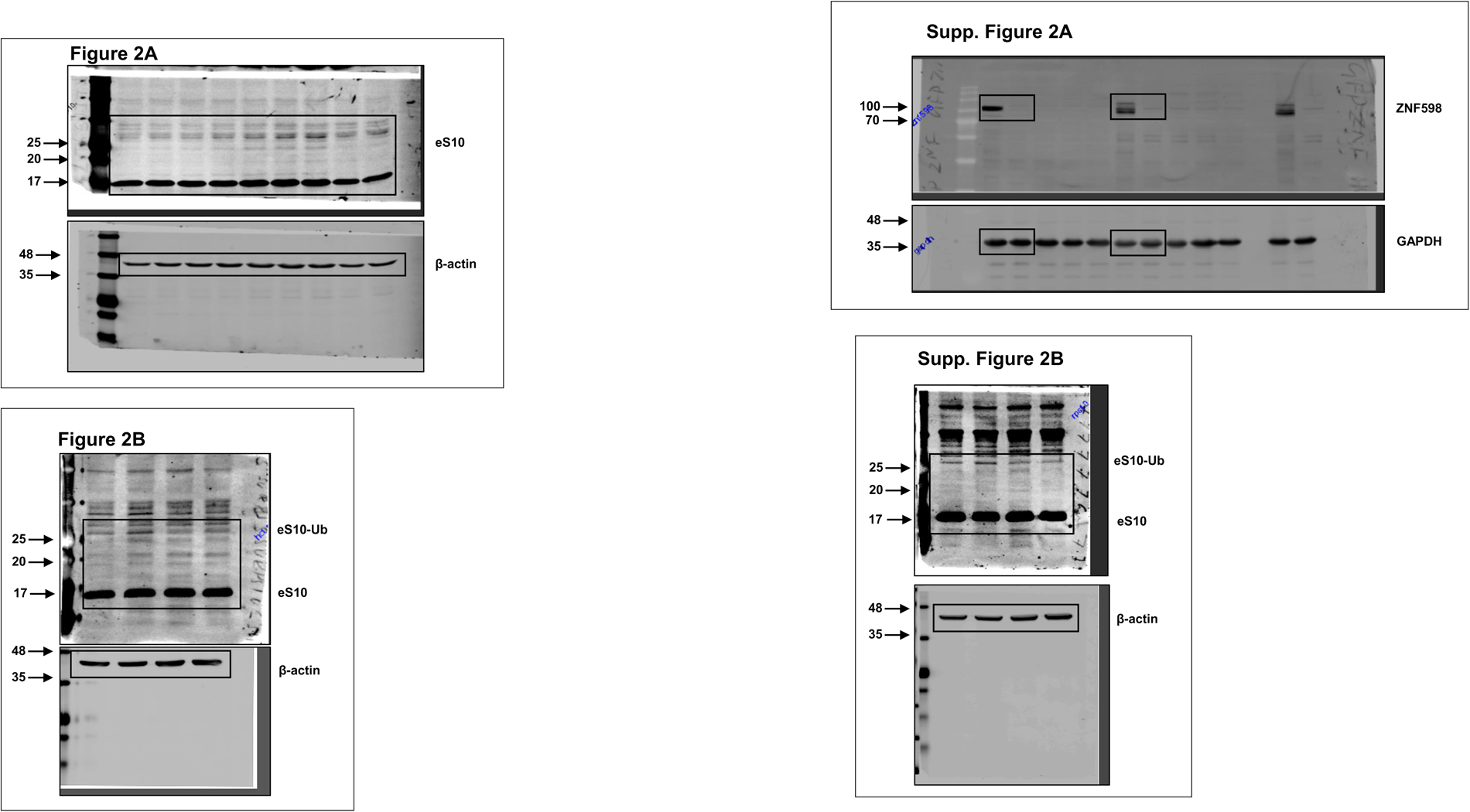
Uncropped images of blots used in Figure 2 and Supp. Figure 2.

**Supplementary Figure 9.**
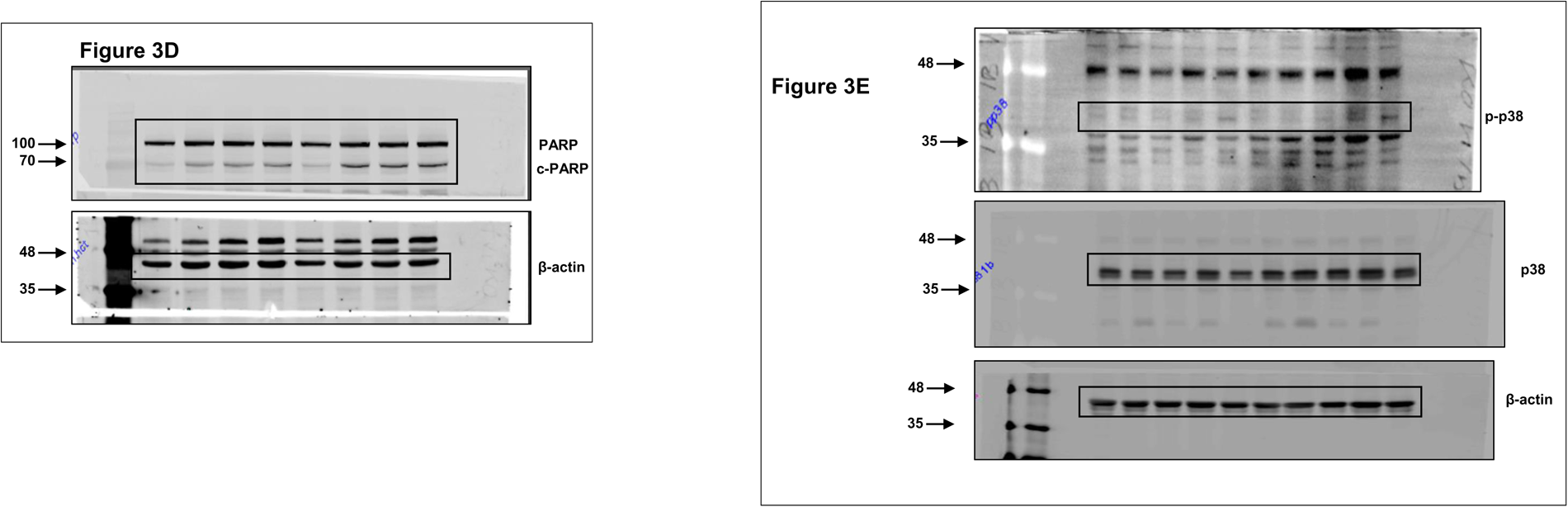
Uncropped images of blots used in Figure 3.

**Supplementary Figure 10.**
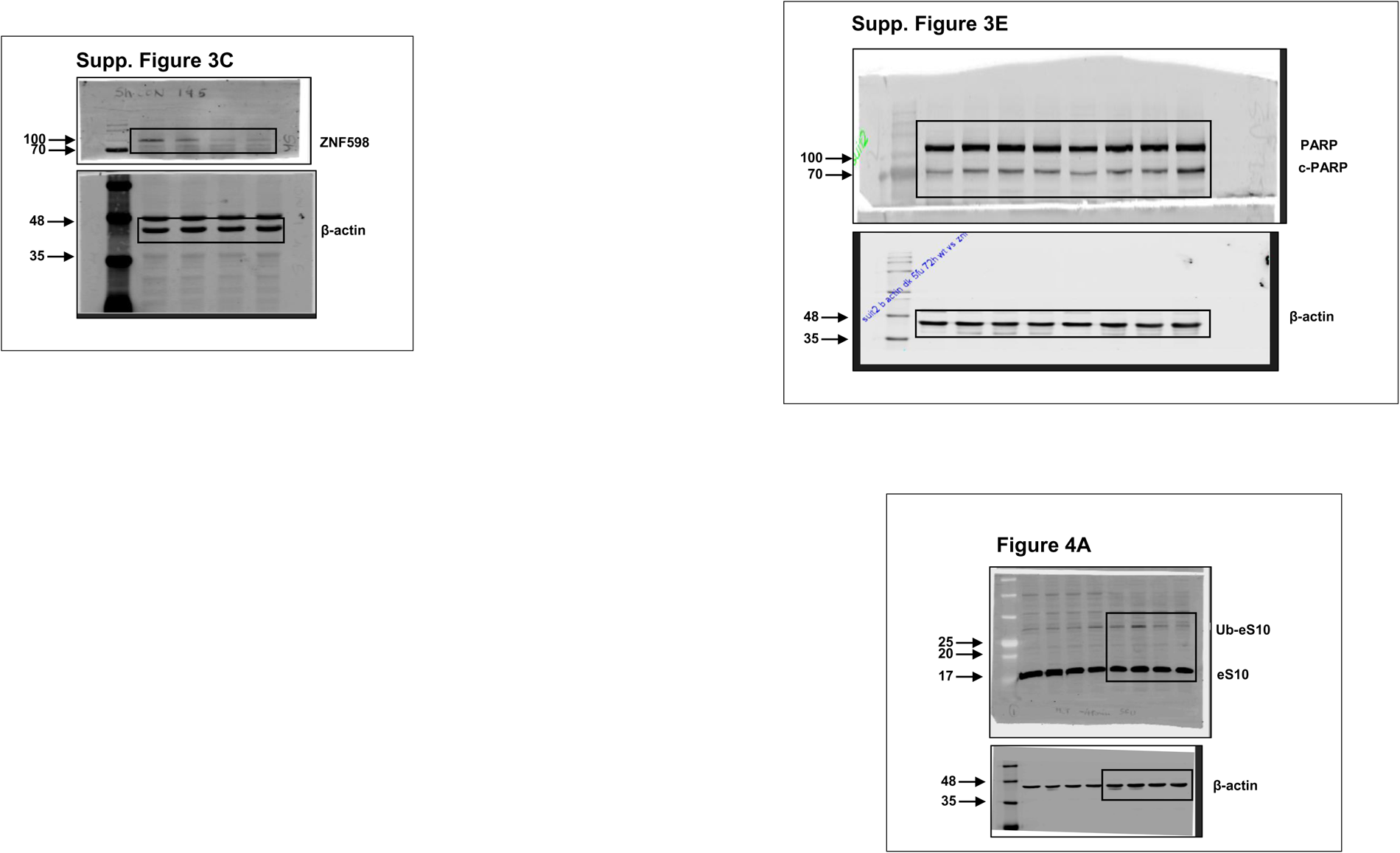
Uncropped images of blots used in Supp. Figure 3 and Figure 4.

**Supplementary Figure 11.**
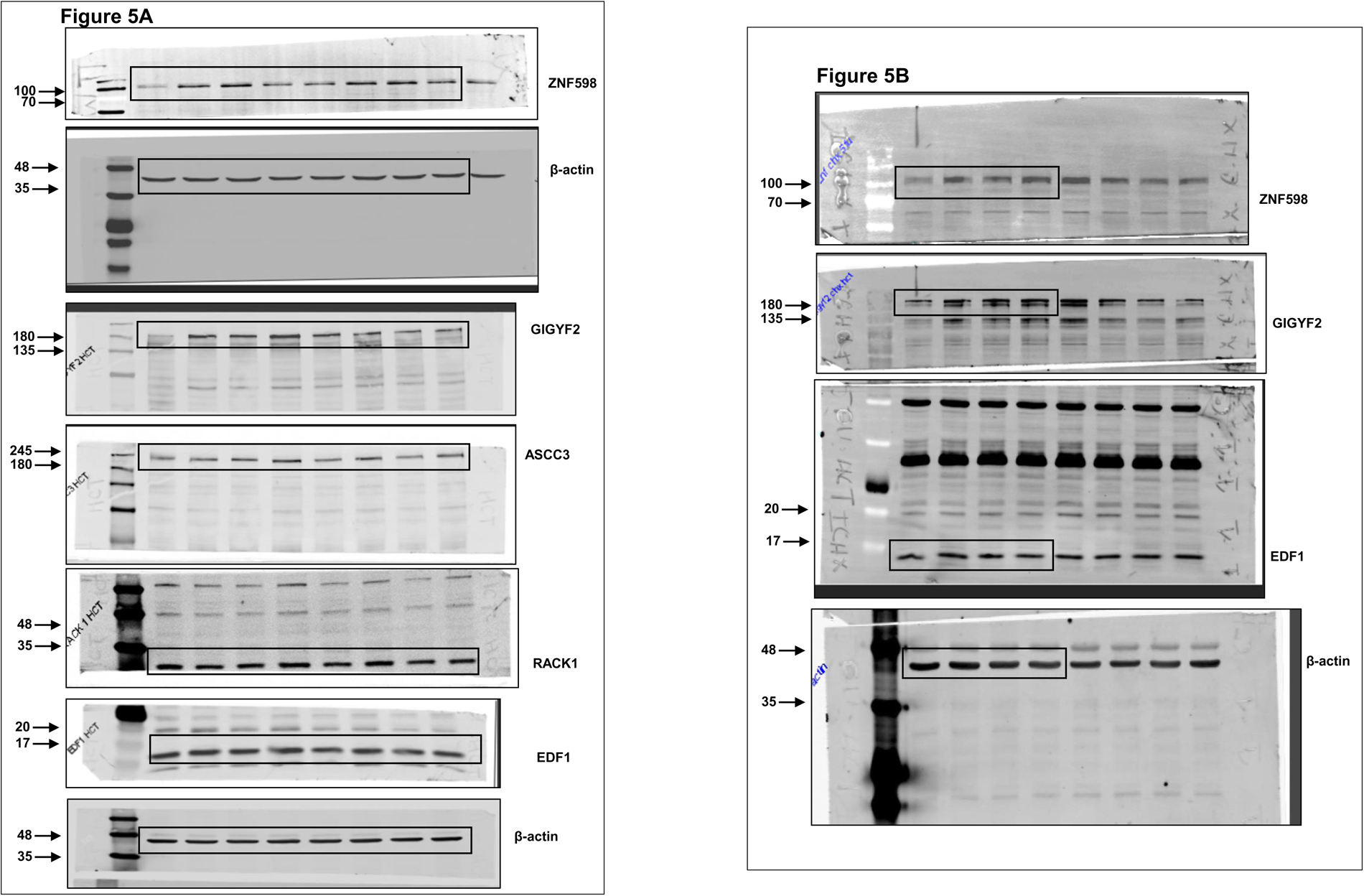
Uncropped images of blots used in Figure 5.

**Supplementary Figure 12.**
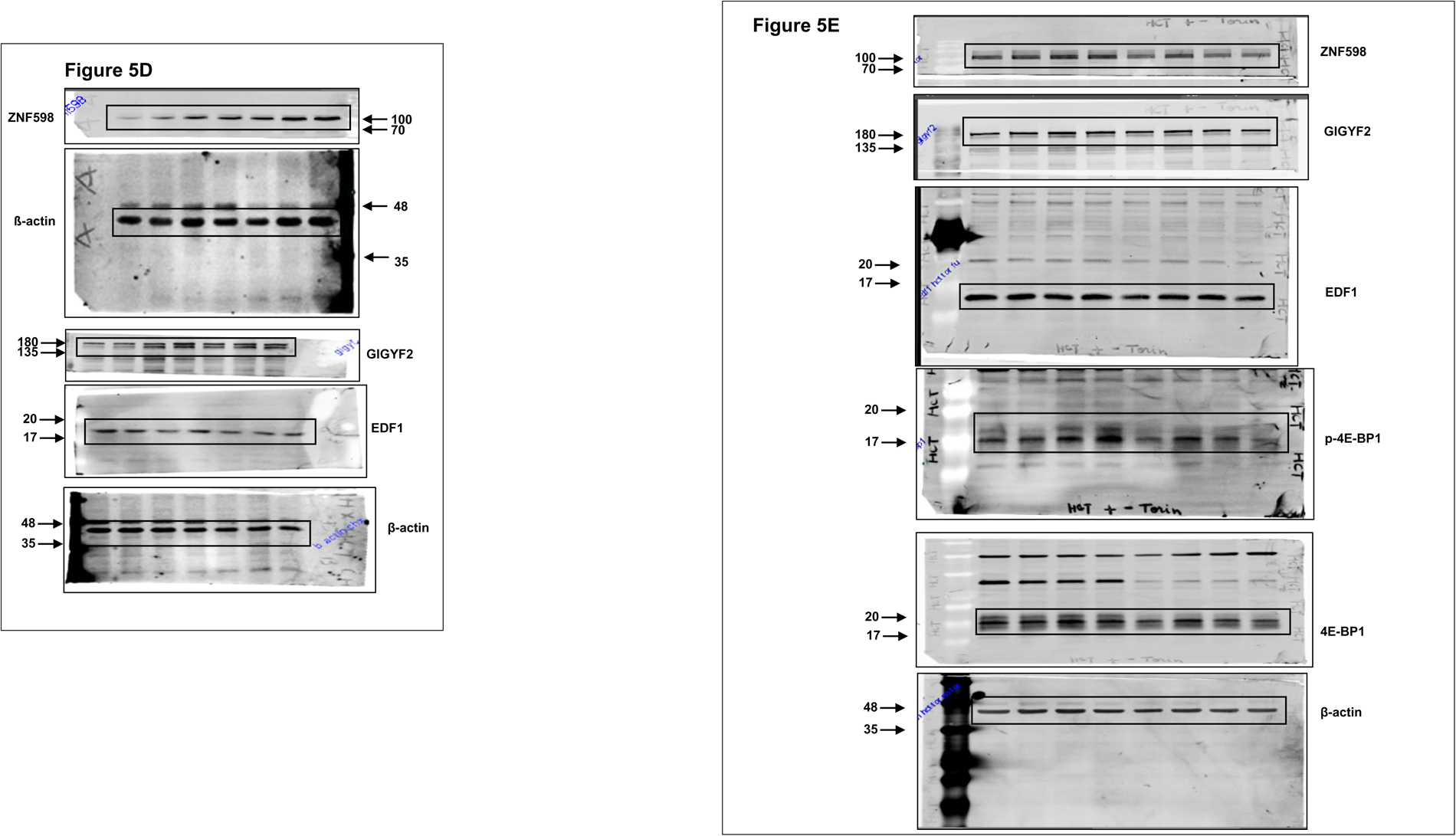

**Supplementary Figure 13.**
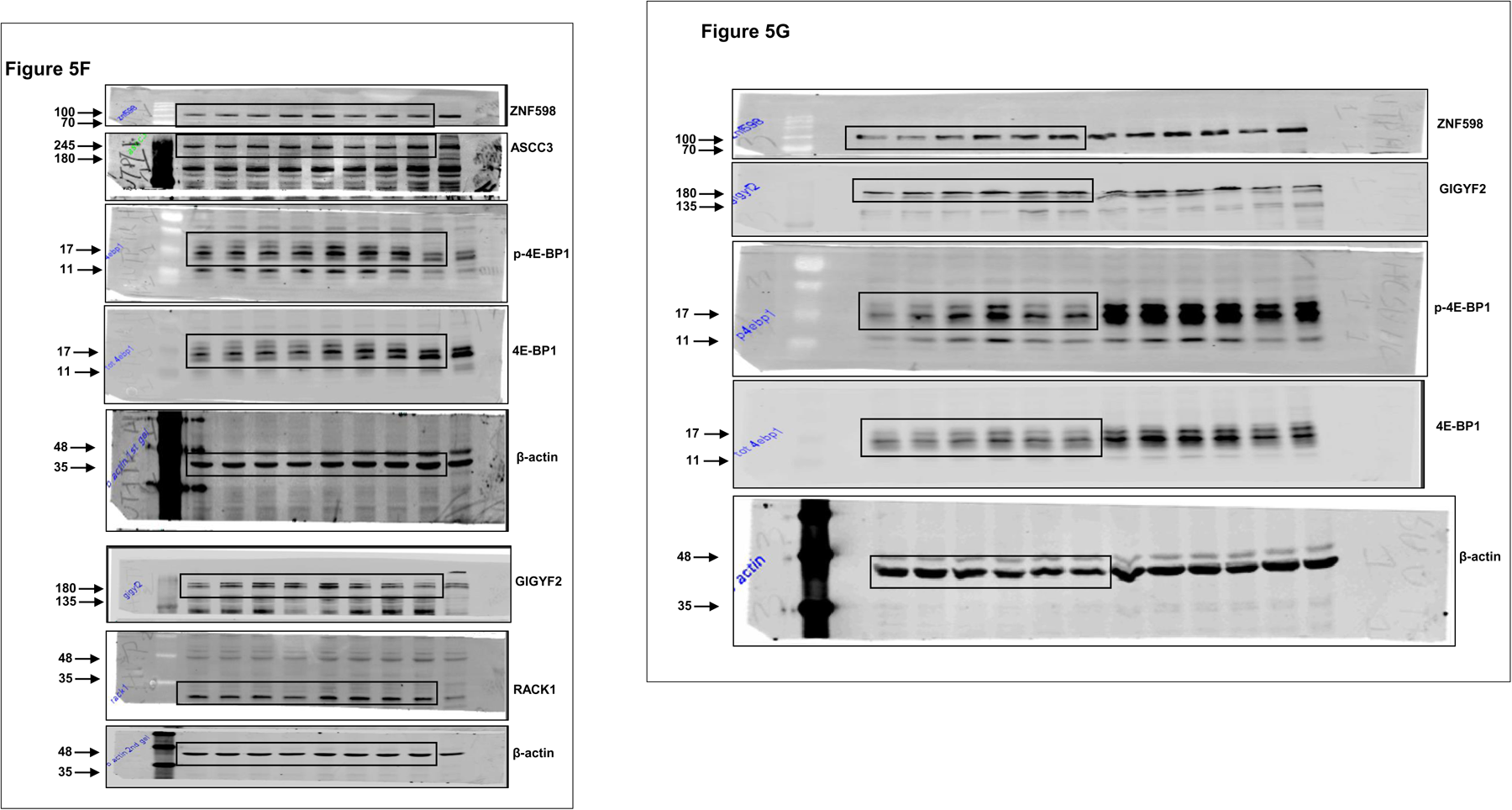

**Supplementary Figure 14.**
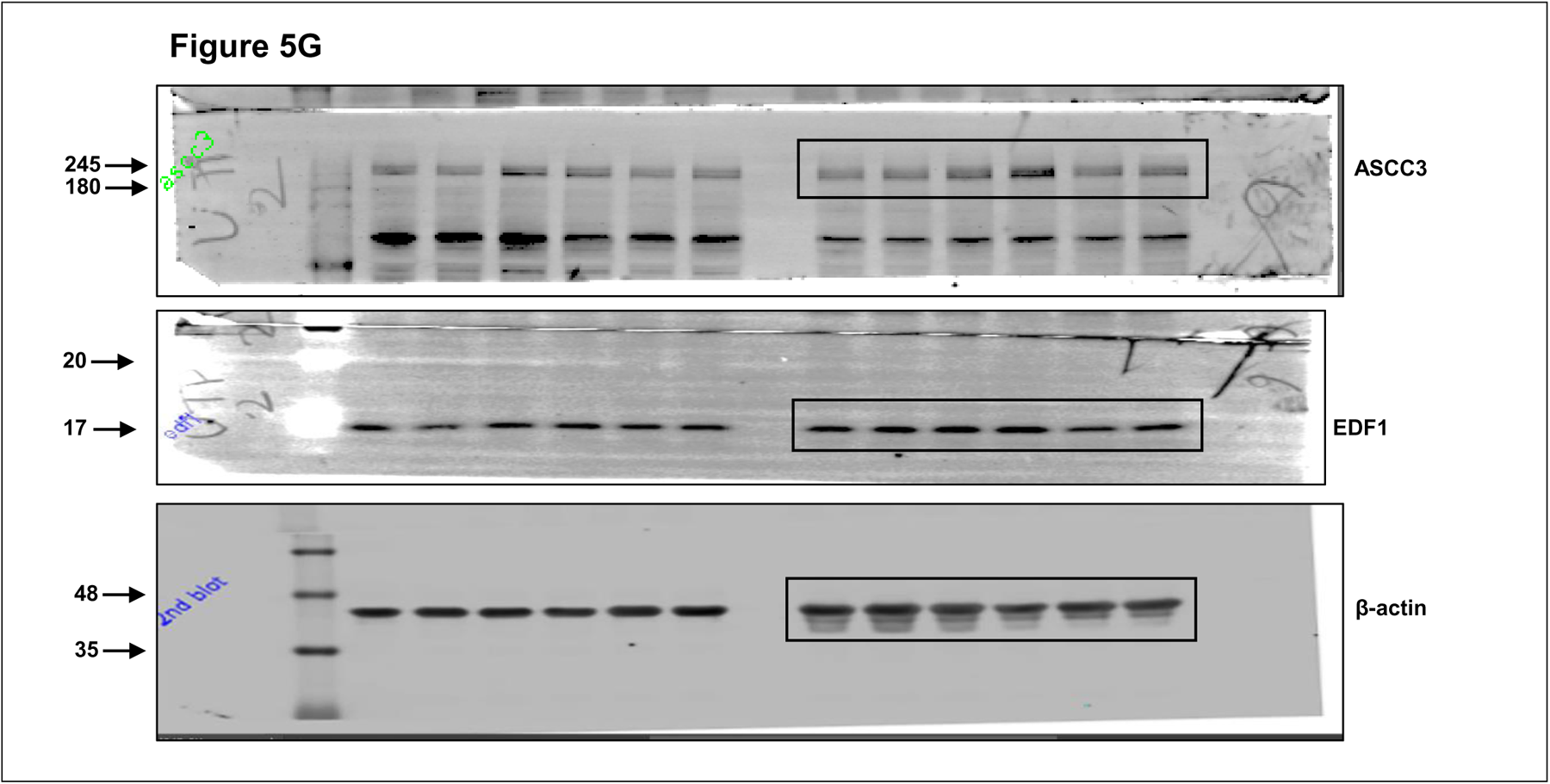

**Supplementary Figure 15.**
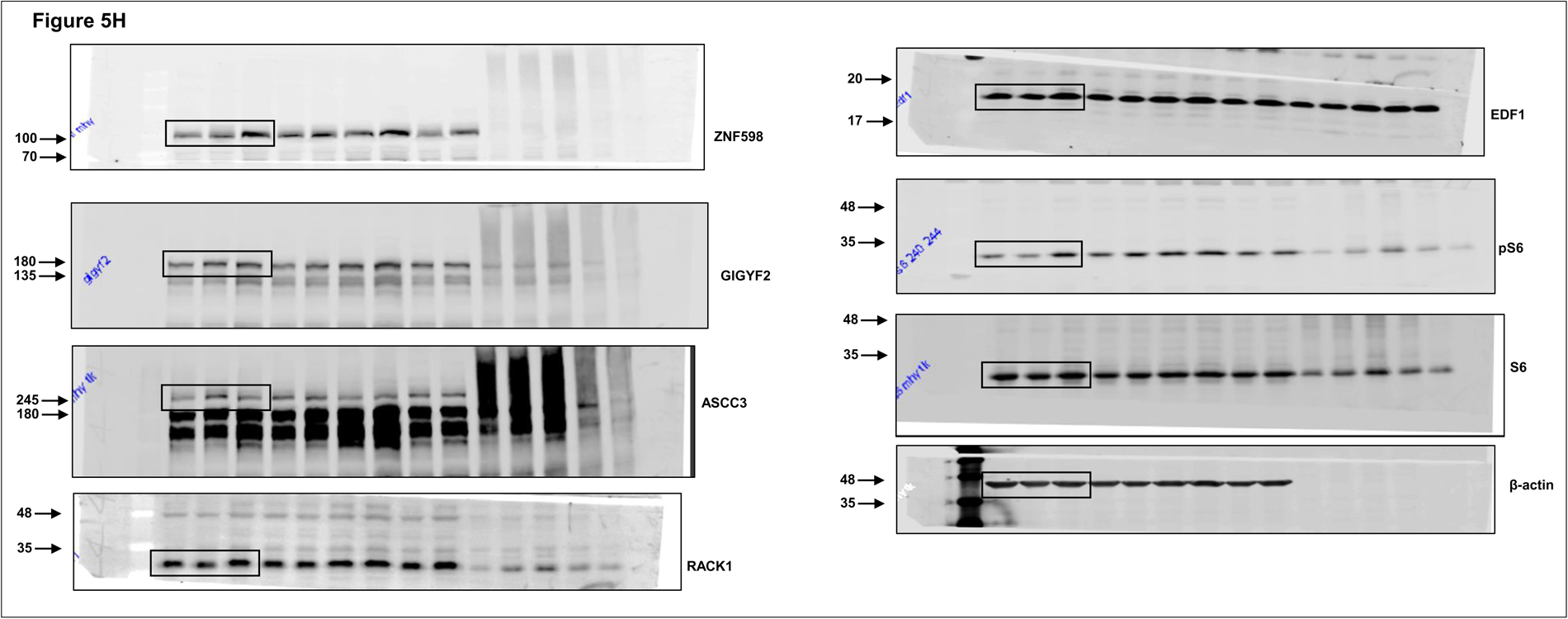

**Supplementary Figure 16.**
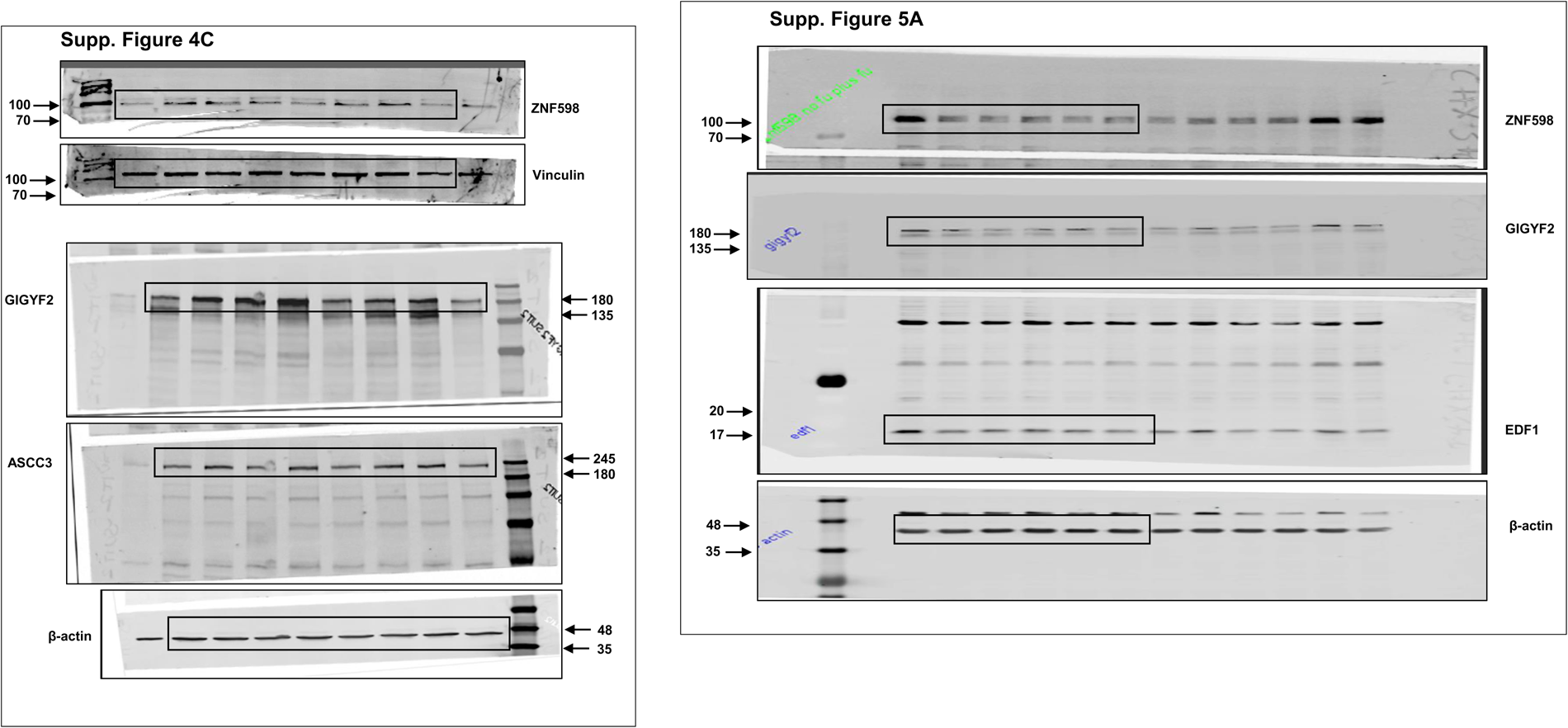
Uncropped images of blots used in Supp. Figure 4 & 5.

**Supplementary Figure 17.**
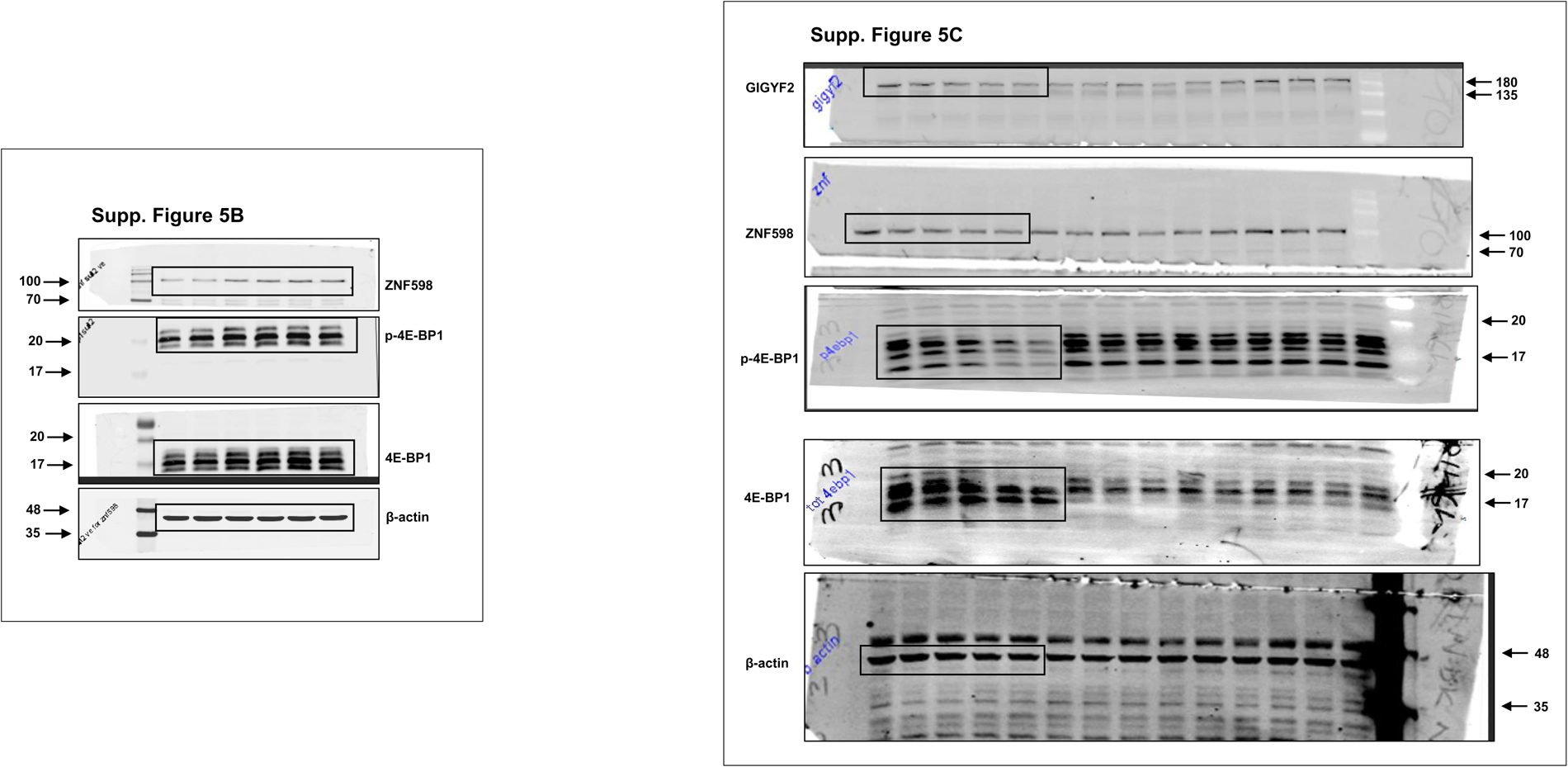
Uncropped images of blots used in Supp. Figure 4 & 5.

**Supplementary Figure 18.**
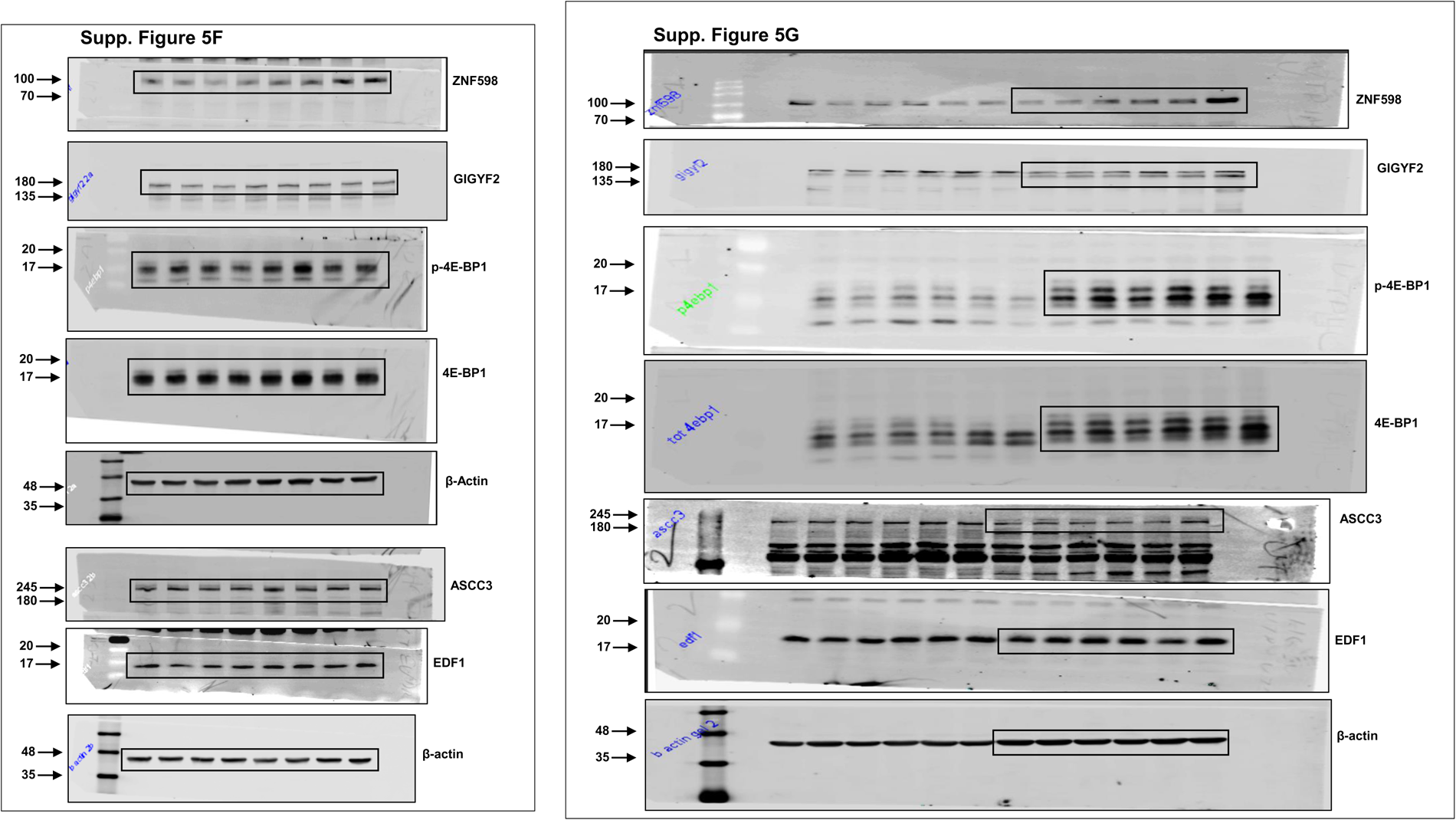
Uncropped images of blots used in Supp. Figure 4 & 5.

